# Nova proteins direct synaptic integration of somatostatin interneurons through activity-dependent alternative splicing

**DOI:** 10.1101/845230

**Authors:** Leena A. Ibrahim, Brie Wamsley, Norah Al-Ghamdi, Nusrath Yusuf, Elaine Sevier, Ariel Hairston, Mia Sherer, Xavier Hubert Jaglin, Qing Xu, Lihua Guo, Alireza Khodadadi-Jamayran, Emilia Favuzzi, Yuan Yuan, Jordane Dimidschstein, Robert Darnell, Gord Fishell

## Abstract

Somatostatin interneurons are the earliest born population of cortical inhibitory cells. They are crucial to support normal brain development and function; however, the mechanisms underlying their integration into nascent cortical circuitry are not well understood. In this study, we begin by demonstrating that the maturation of somatostatin interneurons is activity dependent. We then investigated the relationship between activity, alternative splicing and synapse formation within this population. Specifically, we discovered that the Nova family of RNA-binding proteins are activity-dependent and are essential for the maturation of somatostatin interneurons, as well as their afferent and efferent connectivity. Within this population, Nova2 preferentially mediates the alternative splicing of genes required for axonal formation and synaptic function independently from its effect on gene expression. Hence, our work demonstrates that the Nova family of proteins through alternative splicing are centrally involved in coupling developmental neuronal activity to cortical circuit formation.

## INTRODUCTION

Somatostatin cortical interneurons (SST cINs) constitute ∼30% of all inhibitory interneurons in the cerebral cortex. They are crucial for gating the flow of the sensory, motor, and executive information necessary for the proper function of the mature cortex (Fishell and Rudy, 2011; Kepecs and Fishell, 2014; Tremblay et al., 2016). In particular, Martinotti SST cINs, the most prevalent SST cIN subtype, are present in both the infragranular and supragranular layers of the cortex and extend their axons into Layer 1 (L1) (Rudy et al. 2011; Lim et al. 2018; Ascoli et al. 2008; Nigro, Hashikawa-Yamasaki, and Rudy 2018; Pouchelon et al. 2021) They specifically target the distal dendrites of neighboring excitatory neurons, thus providing the feedback inhibition necessary for modulating dendritic integration (Adler et al., 2019;; Kapfer et al., 2007; Silberberg and Markram, 2007). These roles are dependent upon the ability of SST cINs to form specific synaptic connections with select excitatory and inhibitory cell types during development (Favuzzi et al. 2019).

The mechanisms responsible for generating the precise functional connectivity of SST cINs are poorly understood. Early neuronal activity has emerged as an important factor in directing the maturation of cINs (Wamsley and Fishell, 2017). In addition, recent work has implicated activity as being centrally involved in alternative splicing (Eom et al., 2013; Furlanis and Scheiffele, 2018; Iijima et al., 2011a; Lee et al., 2007; 2009; Mauger et al., 2016; Quesnel-Vallières et al., 2016; Vuong et al., 2016; 2018; Xie and Black, 2001). However, whether these processes are coupled within SST cINs has not been explored.

The Nova family of RNA-binding proteins (Nova1 and Nova2) have been shown to control the splicing and stability of transcripts encoding a variety of neurotransmitter receptors, ion channels, and transmembrane cell adhesion molecules known to affect synaptogenesis and excitability (Dredge and Darnell, 2003; Eom et al., 2013; Saito et al., 2016; 2019; Ule et al., 2005; 2003a; 2006; Yano et al., 2010). Notably both Nova1 and Nova2 are strongly expressed within cINs during periods of synaptogenesis and as such represent promising effectors that may direct the maturation of SST cINs. Here we report that neuronal activity strongly influences efferent SST cIN connectivity. We show that the conditional loss of *Nova1* or *Nova2* phenocopies the effect of dampening activity during circuit assembly, leading to a loss of their efferent inhibitory output. At a molecular level these changes are mediated by a Nova-dependent program, which controls gene expression and alternative splicing of mRNAs encoding for pre- and post-synaptic proteins. Demonstrating a direct link between activity, Nova function, and inhibitory output, increasing activity using NachBac in Nova2 knockouts fails to enhance SST inhibitory output. Conversely, overexpression of *Nova2* within SST cINs marginally increases SST inhibitory output, a phenotype that can be suppressed by damping neuronal activity within these cells. Thus, our work indicates that early activity through a Nova-dependent mechanism is required for the proper establishment of SST cIN connectivity and maturation.

## RESULTS

### Neuronal activity affects the synaptic development of SST cINs

The cortex exhibits a variety of dynamic network activity patterns during cortical synaptogenesis (Allene and Cossart, 2010; Garaschuk et al., 2000; J. W. Yang et al., 2009). These are comprised by both spontaneous and sensory evoked events (Garaschuk et al., 2000; Minlebaev et al., 2011; J.-W. Yang et al., 2012; (Pouchelon et al. 2021; Ibrahim et al. 2021). While inhibitory cortical interneurons (cINs) are recruited by these activities (Cossart, 2011; Le Magueresse and Monyer, 2013), whether this influences somatostatin (SST) cIN development has not been fully established. To address the impact of activity on these cINs, we chose to selectively and cell-autonomously dampen or augment their excitability during the first few weeks of development. This represents a perinatal period in cIN development during nascent circuit formation, where they are robustly forming or losing synaptic contacts (Allene et al., 2008; Minlebaev et al., 2011; J.-W. Yang et al., 2012; J. W. Yang et al., 2009). SST cINs in the primary somatosensory cortex (S1) were targeted using AAV viral injections in *SST^Cre^* mice crossed with a conditional synaptophysin1-eGFP (Syp-eGFP) mouse, which functions as a presynaptic reporter (i.e *SST^cre^;R26R ^LSL-tTa^;Tg-TRE::Syp-eGFP*, Figure1A) (Basaldella et al., 2015; Li et al., 2010; Wamsley et al., 2018). To modulate the activity of SST cINs, these mice were injected at P0 with Cre-dependent AAVs that drive the expression of either KIR2.1 (AAV-Syn-DIO-KIR2.1-P2A-mCherry) or NaChBac (AAV-Syn-DIO-NaChBac-P2A-mCherry) channels coupled to mCherry reporter (Figure 1A). Both channels are voltage-sensitive and have proven to be useful tools to manipulate cellular excitability. The KIR2.1 channel is an inward rectifying potassium channel, which upon overexpression lowers the resting membrane potential towards the reversal potential of K^+^ (∼90mV) (Bortone and Polleux, 2009; De Marco García et al., 2011; Karayannis et al., 2012; Priya et al., 2018; Yu et al., 2004), thus reducing neuronal excitability. The NaChBac channel has an activation threshold that is 15mV more negative than endogenous voltage-gated Na^+^ channels and remains open for 10 times longer (Lin et al., 2010) and therefore augments excitability.

**Figure 1.**
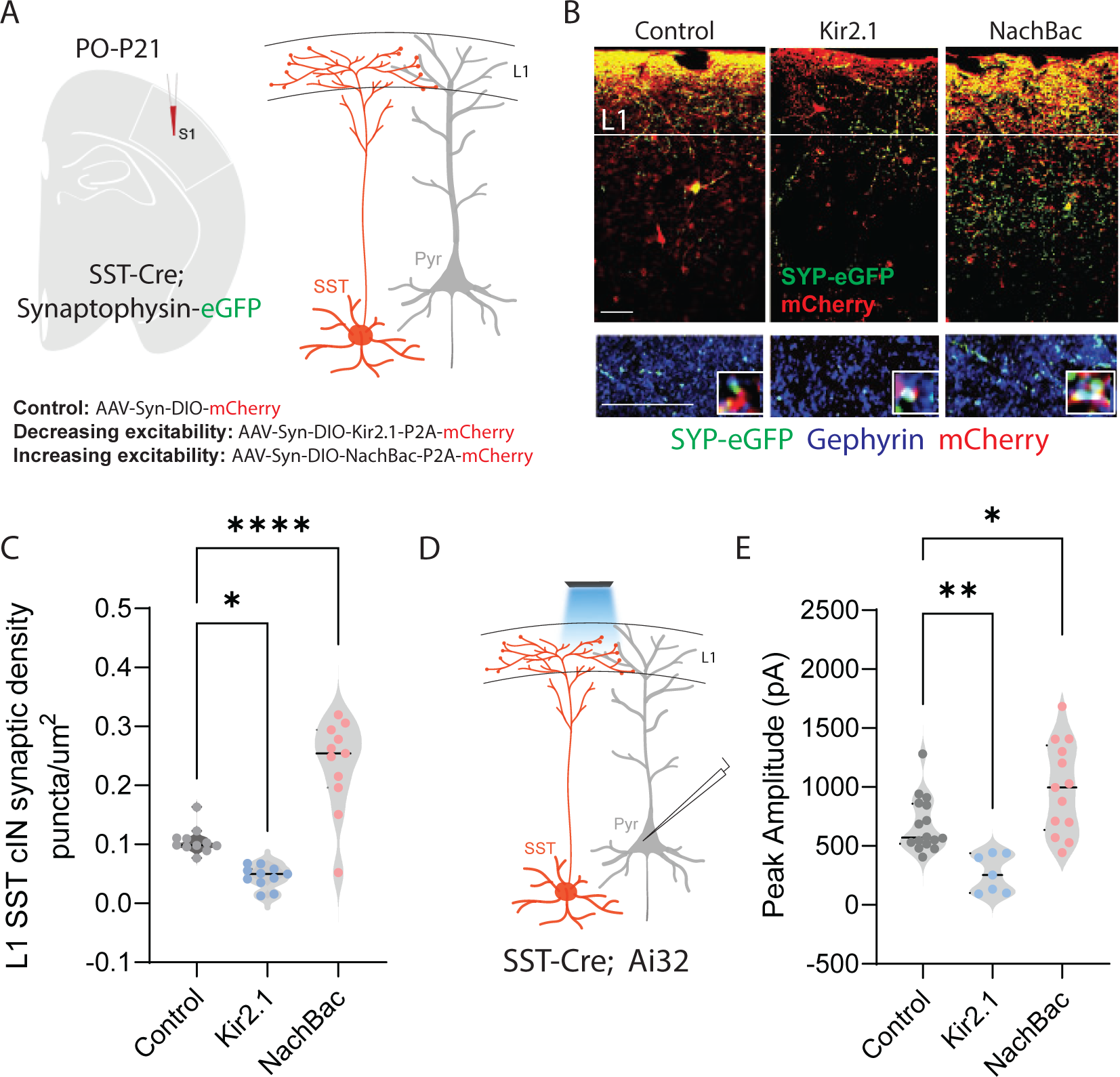
Neuronal activity affects the synaptic development of SST cINs. A) Schematic of genetic alleles (left) and experimental approach (middle), *SSTCre;Syp-eGFP* pups were injected with a conditional virus; either *AAV2/1-Flex-Kir2.1-P2A-mCherry, AAV2/1-Flex-NaChBac-P2A-mCherry,* or Control *AAV2/1-Flex-mCherry* within the S1 cortex at postnatal day 0 (P0). Schematic of the efferent connectivity of SST+ Martinotti cells (right). B) Upper panels: Immunostaining (IHC) of *SSTCre;Syp-eGFP* in layer 1 (L1) of S1 cortex at P21 showing Syp-eGFP (green, anti-GFP) and axons (red, anti-RFP) from *control, Kir2.1, or NaChBac* injected SST+ Marintonotti cINs (scale bar 50um). Lower panels: visualization of Syp-eGFP (green, anti-GFP) and Gephyrin+ puncta (blue, anti-Gephyrin) in L1 SST+ cINs (scale bar 20um). Inset shows a higher magnification image of the puncta overap (Red, mCherry axons). C) Quantification of synaptic puncta (RFP+/GFP+/Gephyrin+ overlap) of control, Kir2.1, and NaChBac expressing SST+ cINs within L1 (n =3-4 mice each, 9 sections each; pVal**= 0.008, ***=0.0001). D) Left, schematic of optogenetic activation of SST neurons using SST-Cre::Ai32 mice injected with either *AAV2/1-Flex-Kir2.1-P2A-mCherry, AAV2/1-Flex-NaChBac-P2A-mCherry,* or Control *AAV2/1-Flex-mCherry* within the S1 cortex at postnatal day 0 (P0) and recording from Pyramidal neurons (clamped at 0mV) in Layer 5 of Primary Somatosensory cortex (S1) at P21. E) Quantification of SST output onto Pyramidal neurons, Peak amplitude of the Inhibitory post synaptic current (IPSC) (Control Peak Amplitude: 663.47±18.7pA, Kir2.1: 185±19.78pA, NachBac: 927.7±28.5pA).

To assess the development of the synaptic efferents of infected SST cINs, we allowed pups to mature until juvenile age. The somatosensory cortex was then subjected to immunohistochemistry (IHC) to visualize pre-synaptic (SST+cIN-mCherry+-Syp-eGFP+) compartments and post synaptic components and subjected to puncta analysis (Figure 1B). We quantified SST efferent synapses identified through the colocalization of the virally mediated mCherry reporter, Syp-eGFP and the postsynaptic marker gephyrin (mCherry+/GFP+/gephyrin+ puncta), as a proxy for synaptic contacts (Ippolito and Eroglu, 2010). In SST cINs, KIR2.1 expression resulted in a significant reduction of L1 SST cIN efferent synaptic puncta in comparison to control cells (0.109±0.009 puncta/um^2^ CTL vs 0.049±0.006 puncta/um^2^ KIR2.1) (Figure 1C). By contrast, the overexpression of NaChBac within SST cINs resulted in a robust increase in L1 synaptic puncta (0.109±0.009 puncta/um^2^ ctl vs 0.218±0.030 puncta/um^2^ NaChBac) (Figure 1C). Additionally, when we optogenetically activated SST neurons and recorded from pyramidal cells (Figure 1D), the inhibitory output was also affected. Kir2.1 expression resulted in a significant reduction in the output of SST+ cINs onto pyramidal cells compared to controls (680±14pA ctl vs 266.5±21pA Kir2.1, Figure 1E). By contrast, the overexpression of NachBac resulted in an increase in the inhibitory output of these cells (680±14pA ctl vs 989.33±28pA, Figure 1E). These results suggest that activity has a profound effect on the density of SST cIN axons, synapses and inhibitory output. Dampening excitability decreases the number of efferent synaptic structures and axonal arbors of SST cINs, while augmenting it increases both.

### Neuronal activity influences alternative splicing and Nova expression within SST cINs

A growing number of studies indicate that activity-dependent alterative splicing (AS) contributes to the regulation of gene expression and the fine-tuning of transcriptional programs related to synaptic refinement (Eom et al., 2013; Iijima et al., 2011b; Fuccillo et al; 2015, Mauger et al., 2016; Quesnel-Vallières et al., 2016; Vuong et al., 2016). This prompted us to test whether neuronal activity itself changes the level of AS within SST cINs during circuit formation, independent of the changes in gene expression. To do so, we used electro-convulsive shock (ECS) during peak synaptogenesis (P8) in mice with genetically-labeled SST cINs (*SST^Cre^;Ai9*). The ECS method generates an acute and reproducible increase in neuronal activity *in vivo* (Guo et al., 2011; Ma et al., 2009), resulting in increased expression of immediate early genes (IEG) such as *Fos*, *Egr1*, *Npas4* and *Arc* (Figure S1A-F), analogous to that observed with KCl treatment *in vitro* but with the added advantages of being *in vivo* and transient. Two to three hours following ECS, we isolated SST cINs from the S1 cortex of *SST^Cre^;Ai9* animals using fluorescence activated cell sorting (FACS) (Figure 2A and Figure S2A). Sorted SST cINs were used to prepare cDNA libraries that were subsequently sequenced in order to investigate changes in AS (spliced exon: SE, mutually exclusive exons: MXE, retained intron: RI, alternative 5’ splice site: A5, alternative 3’ splice site: A3) (Figure 2A right). We found 312 transcripts differentially spliced between sham/control and ECS (FDR<0.05, |Δψ|≥0.1 threshold), comprised by 139 SE events (57 excluded and 82 included exons), 66 RI events (13 excluded introns and 53 included), 31 MXE events (29 excluded and 26 included exons), 13 A5 events (1 excluded and 12 included exons), and 39 A3 (23 excluded and 16 included exons) (Figure 2B).

**Figure 2.**
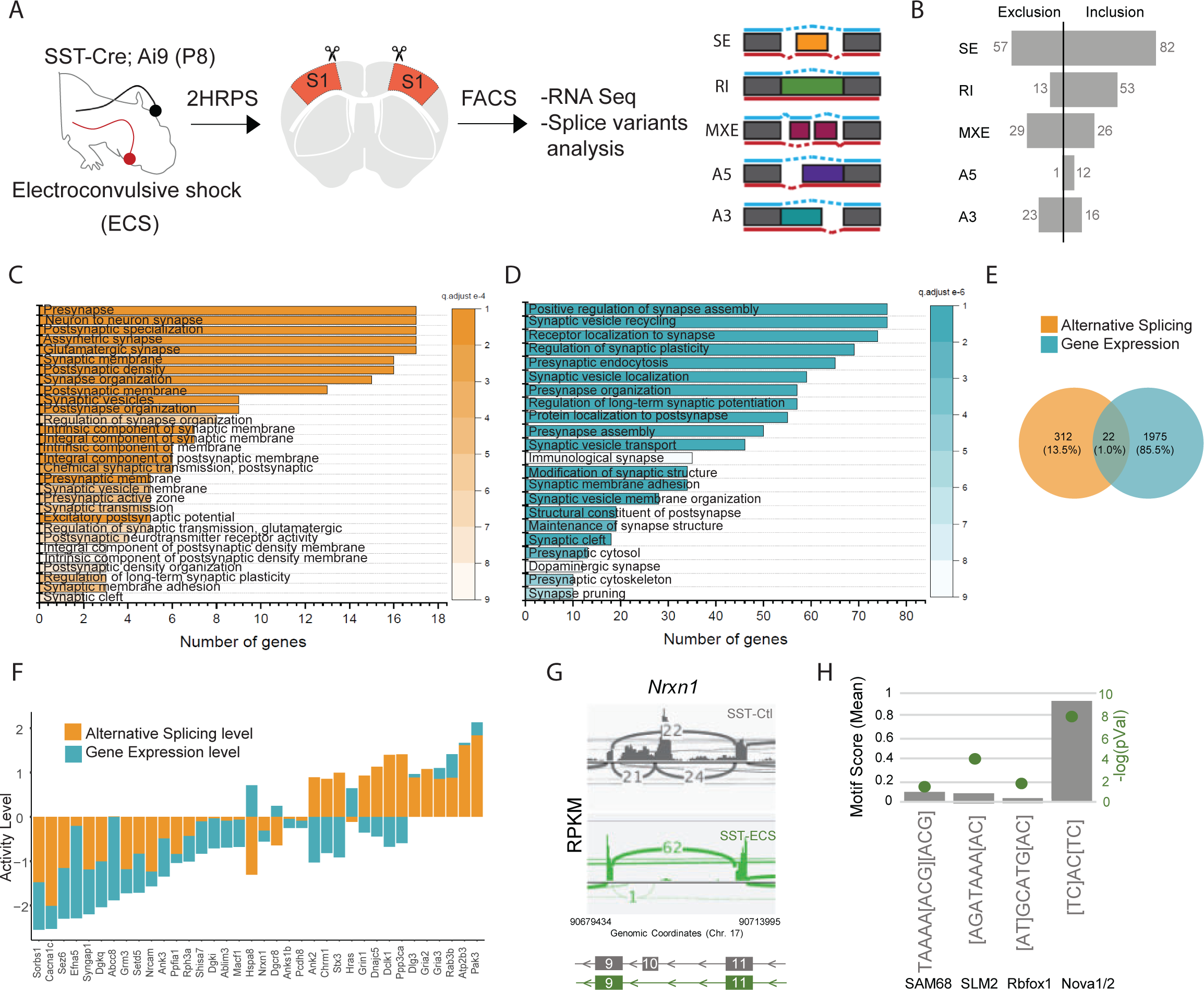
Neuronal activity influences alternative splicing and Nova expression within SST cINs. A) Schematic of experimental approach: Postnatal day 8 (P8) *SSTCre;Ai9* pups were subjected to electroconvulsive shock (ECS) (left). Following 2-3 hours the S1 cortex was isolated and SST+ cINs were FACS purified (middle). SST+ cINs were then prepared for RNAseq to assess changes in gene expression and alternative splicing. Splicing changes are divided into the major alternative structural motifs: single exon, SE, retained intron, RI, mutually exclusive exons, MXE, alternative 5’ splice site, A5, alternative 3’ splice site, A3 (right). B) Histogram of the magnitude of activity-dependent splicing changes within SST+ cINs subjected to ECS compared to sham SST+ cINs (FDR <0.5, fold <0.1>), depicting 139 differential spliced SE (82 SE included, 57 SE excluded), 66 differential spliced RI (53 RI included, 13 RI excluded), 55 differential spliced MXE (26 included, 29 excluded), 13 differential spliced A5 (12 included, 1 excluded), 39 differential spliced A3 (16 included, 23 excluded). C) Gene Ontology (GO) analysis of differentially alternatively spliced (AS) genes (Orange color) under synaptic categories. D) Gene Ontology (GO) analysis of differentially expressed genes (GE) (Teal color) under synaptic categories. E) Overlap of all differentially expressed (teal) and differentially spliced (orange) genes under ECS vs control conditions. Below: overlap for synaptic gene only. F) Comparison of activity level of the overlapped synaptic genes (genes that have both AS and GE changes). Activity level is calculated by considering both FC and pvalue. G) Sashimi plot illustrating *Nxrn1* exon 10 exclusion in activity-induced SST cINs in green (bottom) compared to sham SST cINs in grey (top). Reads per kilobase of transcripts (RPKM) gives the count of the number of transcripts for a specfic isoform. H) Histogram of the average motif enrichment score of known activity-regulated splicing factors KHDRBS1 (Sam68), KHDRBS2 (SLM2), Rbfox1 and Nova1/2 (right). Green dots represent -log10 adjusted p value (right Y-axis) for motif enrichment scores, only significant enrichment shown.

Utilizing the SST cIN transcriptome as a reference, we performed gene ontology (GO) analysis to ask if the genes subject to alternative splicing (AS) were enriched for specific functional categories within these neurons. GO analysis of the genes that underwent activity-dependent AS belong to specific ontological categories, such as synapse maturation, synaptic transmission, and axonal growth (Figure 2C and S2B). In addition, and as expected we also observed activity dependent changes in gene expression (Figure 2D). However, the overlap between the genes subjected to AS vs GE was only ∼1.2% of all differentially expressed genes (Figure 1E). When we compared the overlapped genes (those that underwent both GE and AS changes), we observed that for many synaptic genes, the level of AS changes was higher than the changes of the same genes at the transcript level (Figure 1F). This suggests that the changes observed in synaptic genes for AS are independent of their changes in transcription level. For example, we observed and validated (Table S3) that within activity stimulated SST cINs the *Nrxn1* mRNAs exclude exon 10. Notably, this exon lies within a laminin-protein coding domain important for the cell adhesion properties of Nrxn1 at the synapse (Figure 2G, control, grey vs ECS, green). In contrast, while we also observed GE changes in Nrxn1, the level of AS change was higher.

We next asked whether the activity-dependent AS genes formed a protein-protein interacting network (PPI) based on previously established direct protein interactions *in vivo* (Rossin et al., 2011). Notably, the genes subjected to activity-dependent AS within SST cINs form highly connected networks illustrating they likely function together to support pre-synaptic vesicle function (Figure S2C, pink), post-synaptic organization and receptor-associated synaptic components (Figure S2C, blue) (pVal<0.0009, 1000 permutations). These genes among others include: *Hspa8, Nrxn1, Syngap1, Cacna1c, Ppp3ca,* and *Grin1* (Figure S2C). These results indicate that augmenting activity within SST cINs during nascent circuit development robustly increases AS events and most of the spliced mRNAs are genes specifically related to axonal development and synaptic transmission (Figure S2C, pink and blue, respectively).

We next sought to identify RNA binding proteins (RNABPs) that could mediate activity dependent AS events within SST cINs. To do so, we utilized the RNAseq experiments described above (control vs. ECS) to perform a motif enrichment analysis that utilizes position probability matrices of binding motifs from 102 RNABPs (i.e. PTBP1/2, FUS, ELAVL4, SRRM4, Rbfox1, FMR1, Nova1, Nova2) (Liu et al., 2017; Park et al., 2016; Y. Yang et al., 2016). Previous HITS-CLIP analysis has revealed that Nova1 and Nova2 share an almost identical RNA-binding domain (YCAY) (Licatalosi et al., 2008; Ule et al., 2006; 2003b; Yuan et al., 2018a). Strikingly, the Nova-binding motif was found to be significantly enriched within activity-dependent targets and at a higher frequency than other neuronal splice factors (i.e. Sam68 (KHDRBS1), SLM2 (KHDRBS2), and Rbfox1) (pVal<0.0001) (Figure 2H). This finding implicates Nova proteins as playing a fundamental role in directing SST cIN activity-dependent AS.

### Neuronal activity during cortical development influences the expression and localization of Nova proteins in SST cINs

We next examined the expression of Nova1 and Nova2 within SST cINs across development and whether their expression is affected by changes in neuronal activity. Utilizing IHC and genetic fate mapping, we observed that the expression of the Nova family (Nova1 and Nova2) proteins begins within cIN populations soon after they become postmitotic and expressed in 100% of SST and PV cINs by adulthood (Figure S3A, B). For comparison we also examined Nova expression in 5HT3aR cINs (Figure S3B) within this same region. To specifically examine the expression of Nova1 and Nova2 during SST cIN synaptogenesis, we performed quantitative-PCR (qPCR) on FACS isolated cINs from the S1 cortex of *Tg-Lhx6::eGFP* mice at P2, P8, and P15. The *Tg-Lhx6::eGFP* mice express eGFP in both SST and Parvalbumin (PV) cINs (medial ganglionic eminence derived cINs) soon after they become postmitotic. We found that both *Nova1* and *Nova2* are expressed within all SST and PV cINs across the first two weeks of postnatal development, coinciding with nascent circuit development (Figure S3C). Taken together, we find that both Nova1 and Nova2 proteins are highly expressed in all SST cINs during circuit integration and may therefore control integral aspects of their development through activity-dependent alternative splicing.

We next investigated whether Nova1 and Nova2 are activity-regulated within SST cINs by examining both their expression and localization during the peak of nascent circuit integration. To do so, we subjected *SST^Cre^;Ai9* mice to ECS during synaptogenesis, similar to what was done in Fig 2. Next, we performed both RNA Sequencing and qPCR for *Nova1* and *Nova2* within FAC sorted cINs from S1 cortex of either control or ECS-treated animals. Following 2 hours post seizure-induction (2 HRPS), we found that the mRNA expression levels of both *Nova1* and *Nova2* were increased in ECS-treated SST cINs compared to controls (*Nova1:* 28.4±5.59 ECS vs 7.61±0.25 control; *Nova2* pVal=0.002: 10.32±1.80 ECS vs 6.59±0.28 control pVal=0.005, Figure S3E and Figure 3A). Next, we probed Nova1 and Nova2 protein levels using western blot (WB) of sorted SST cINs from S1 cortex. Consistent with an activity-mediated upregulation in *Nova* expression, we found a significant increase in both Nova1 and Nova2 protein levels (0.824±0.0412 pixel density (pd) Nova1 control vs 5.62±0.969 pd Nova1 2HRPS, pVal= 0.038 and 0.997±0.409 pd Nova2 control vs 5.7±0.582 pixel density Nova2 2HRPS, pVal= 0.022, pixel densities normalized to *ß-*Actin) (Figure 3B). Thus, these results confirm an increase in *Nova* mRNA and Nova protein expression in SST cINs following an acute increase in neuronal activity.

**Figure 3.**
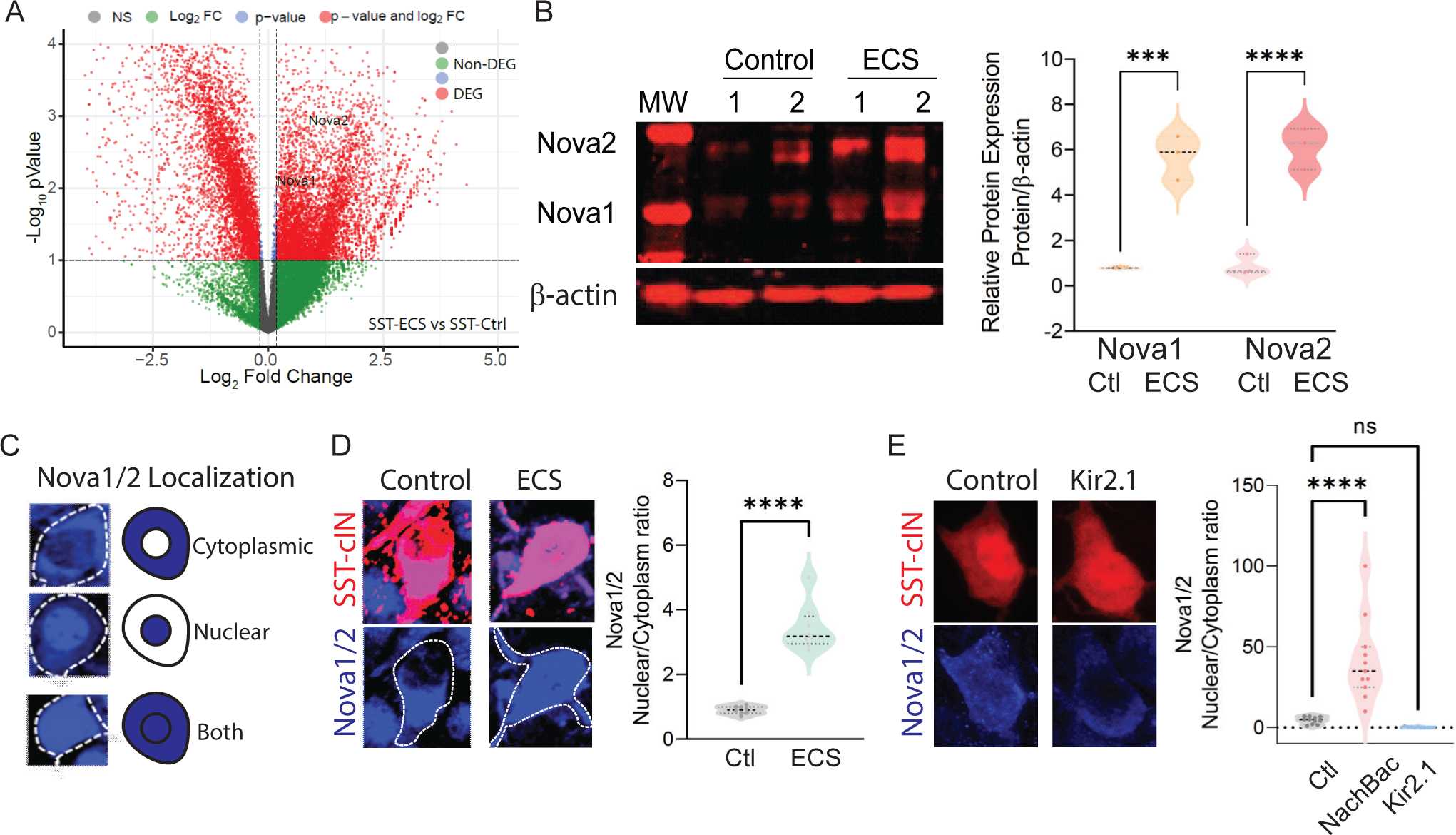
Neuronal activity during cortical development influences the expression and localization of Nova proteins in SST cINs. A) Volcano plot of RNA seq data showing Nova1 and Nova2 upregulation in SST-cINs in ECS vs control. B) Upper panel, western blot showing Nova1 and Nova2 protein expression in control (lanes 2 and 3) versus ECS induced SST cINs (lanes 4 and 5). Lower panel, same western blot showing expression of b-actin across lanes. Right, Quantification of the western blot data. Nova1 and Nova2 protein expression relative to β-actin in control versus ECS induced SST cINs (n=3 mice, S1 cortex only; *pVal=0.038, Nova1; *pVal=0.022, Nova2). C) Representative scoring criteria for Nova1/2 localization within SST cINs : IHC of Nova1/2 (blue, anti-Nova1/2) in selective SST+ cINS exemplifying the Nova1/2 expression in: cytoplasm only (top), nucleus only (middle) and in both cytoplasm and nucleus (bottom). D) Left, representative images of Nova1/2 expression (blue) in SST cINs (red) under normal versus ECS. Right, Quantification of the ratio of nuclear to cytoplasmic localization of Nova1/2 in SST+ cINs of control animals (grey) and ECS animals (green) (n=3 mice, S1 cortex; **pVal= 0.001). E) Left, representative images of Nova1/2 expression (blue) in SST cINs (red) using control mCherry versus Kir2.1-mCherry virus injection. Right, Quantification of the ratio of nuclear to cytoplasmic localization of Nova1/2 in SST+ cINs of control *AAV2/1-Syn-DIO-mCherry* (grey) versus *AAV2/1-Syn-DIO-NaChBac-P2A-mCherry* (pink) versus *AAV2/1-Syn-DIO-Kir2.1-P2A-mCherry* (blue) injected animals. (n=11 mice, S1 cortex, ∼30 cells each; ***pVal=0.0004, NachBac; ***pVal=0.0001, KIR2.1).

Following seizure activity Nova proteins have been shown to translocate into the nucleus within excitatory neurons (Eom et al., 2013). We next sought to explore whether manipulating activity also influences intracellular localization of Nova proteins within SST cINs (Figure 3C). We hypothesized that an activity-mediated increase would direct Nova proteins to the nucleus. We therefore analyzed the ratio of Nova expression within the nucleus versus the cytoplasm of SST cINs following ECS (Figure 3D) or after constitutive activity-modulation across the first postnatal month (DIO:AAV injections of KIR2.1 or NaChBac into *SST^cre^* animals at P0) (Figure 3E). First, we examined Nova localization using IHC 2 HRPS following ECS within the S1 cortex of P8 *SST^Cre^;Ai9* mice (Figure 3C-D). We quantified the proportions of SST cINs that express Nova proteins most prominently within the nucleus versus the cytoplasm by taking multi-Z-stack images of SST cINs and utilizing DAPI to demark the nuclear boundary. We found that Nova localization was observed in three basic patterns in SST cINs: restricted to the cytoplasm, nuclear restricted, or a combination of both nuclear and cytoplasmic expression (Figure 3C). At P8, in the majority of SST cINs, Nova is either restricted to the cytoplasm or expressed in both the nucleus and cytoplasm (Figure 3D). Following an acute increase in activity (ECS), we found a significant increase in the ratio of nuclear to cytoplasmic Nova protein within SST cINs (0.948±0.055 ratio in control versus 3.65±0.465 ratio in ECS, pVal= 0.001, Figure 3D right). Next, by utilizing the same analytical approach, we examined Nova protein localization in S1 of P21 mice that express either KIR2.1 or NaChBac along with an mCherry reporter (Figure 3E). We found a substantial increase in the ratio of SST cINs that localized Nova protein in the nucleus compared to the cytoplasm in NaChBac-expressing expressing SST cINs compared to control cells (4.75±0.678 ratio control vs 40.91±8.41 ratio NaChBac, pVal= 0.0004) (Figure 3E right). In contrast, we found a small decrease in the ratio of nuclear to cytoplasmic Nova protein within SST cINs injected with KIR2.1 (4.75±0.678 ratio control vs 0.265±0.104 ratio KIR2.1, pVal= <0.0001) (Figure 3E right). Most strikingly, we also observed more than half of SST cINs subjected to KIR2.1 either do not express Nova or have substantially reduced levels of Nova protein expression (Figure S3F-G), suggesting that normal levels of activity are needed for maintaining Nova protein expression in the cell. Altogether these data indicate that during synaptogenesis Nova protein expression and localization within SST cINs is strongly modulated by acute or persistent changes in activity.

### Nova1 and Nova2 control distinct AS networks within SST cINs

To address whether Nova1 and Nova2 differentially affect connectivity and maturation, we asked what AS networks they control within SST cINs during development. Given that they share a very similar RNA-binding motif and are found associated with one another in vivo, they were thought to function cooperatively (Licatalosi et al., 2008; Racca et al., 2010; Yuan et al., 2018b). However recently, it has been shown that in addition to their synergistic roles, Nova1 and Nova2 proteins each control distinct AS gene networks (Saito et al., 2019; 2016). We thus chose to examine changes in AS within SST cINs in Nova1, Nova2 or Nova1/2 compound conditional knockout mice (Saito et al., 2019; Yuan et al., 2018b). Using FAC sorting, we isolated SST cINs from *SST^Cre^;Nova1^F/F^* or *SST^Cre^*; *;Nova2^F/F^* or *SST^Cre^;Nova1^F/F^*; *Nova2^F/F^* mice on an Ai9 reporter background (referred to henceforth at *SST-Nova1*, *SST-Nova2* and *SST-dKO,* respectively) at P8 (Figure 4A). We prepared cDNA libraries from FAC sorted SST cINs, performed RNA sequencing and assessed AS changes between control SST cINs versus each of these mutant alleles. Compared to wild type controls, *Nova1* loss resulted in 124 altered AS events (81 excluded and 43 included), *Nova2* loss lead to 339 altered AS events (217 excluded and 122 included) and double mutants exhibited 270 altered AS events (162 excluded and 108 included) (FDR < 0.05) (Figure 4B). Notably, within SST cINs, the loss of *Nova2* results in the largest number of changes in mRNA splicing events compared to compound loss of either *Nova1* or both *Nova* genes. Interestingly, the loss of *Nova1*, *Nova2* or *N1/2-dKO* also resulted in significant gene expression changes within SST-cIN (Figure S4A-F). However, similar to what we observed with ECS, many of the genes that were subjected to AS (Figure 4C) were independent from the genes that underwent changes in gene expression (Figure 4D). Amongst the common genes (between AS and GE), many synaptic genes showed higher levels of AS changes compared to gene expression changes (Figure 4D).

We next assessed the overlap of changes in AS events observed within each mutant (Figure S5A-C). We found the number of alterations in *SST-Nova2* AS events that overlap with *SST-dKO* is almost three times higher than that observed when comparing the overlap between *SST-dKO* and *SST-Nova1* (i.e. 62 altered *SST-Nova2* AS events coincided with the 162 observed in *SST-dKO* versus an overlap of only 25 AS events that were altered in *SST-Nova1* mutants, Figure S5B, C). By contrast, less than 15% of the altered *SST-Nova1* AS genes overlap with changes observed in *SST-Nova2* mutants (i.e. only 28 of the 217 *SST-Nova2* events were altered in *SST-Nova1* mutants) (Figure S5A, B). Interestingly, *SST-dKO* mutants exhibited less altered splicing events than the single *SST-Nova2* mutant, suggesting that some inclusion and exclusion AS events are antagonistically directed by Nova1 and Nova2. Additionally, we performed a correlation analysis within and between Nova1, Nova2 and Nova1/2-dKO to assess whether the type of splicing events is correlated. Similar to above, we observed a higher correlation in exclusion events between Nova2 and Nova1/2-dKO, compared to Nova1 and Nova1/2-dKO (Figure S5D).

To infer their specific biological functions, we performed GO analysis on the altered AS events from each mutant and then asked whether the affected AS events form direct PPI networks. GO analysis of the *SST-Nova1* targets did not result in any significant enrichment of specific functional categories (below an FDR of 0.05) however it did organize genes into categories such as RNA binding, ion binding, and catalytic activity (Figure S4G). *SST-Nova1* AS genes formed a relatively indistinct small sparse PPI network (pVal<0.09) representing vesicle-transport and nucleic-acid binding pathways (Figure S4J, pink shaded). In contrast, *SST-Nova2* and *SST-dKO* AS genes organized into several shared significant GO categories such as neuron projection, axon, cell-cell junction, and synaptic function (FDR <0.05) (Figure S4H, I). *SST-dKO* AS genes also organized into some unique categories, which were involved in postsynaptic specialization, dendrite, and synaptic vesicle membrane (FDR <0.05) (Figure S4H). We next asked if the AS genes affected in *SST-Nova2* and *SST-dKO* were predicted to function together in a PPI network representing specific biological processes (Figure S4K-L). Perhaps not surprisingly both formed highly connected significant PPI networks (pVal < 0.0009, 1000 permutations) representing multiple pathways for vesicle-transport, pre- and post-synaptic function and organization, as well as Ca^2+^ signaling (Figure S4K, L, pink and green respectively). Interestingly, the PPI network for *SST-Nova2* uniquely includes numerous glutamate receptors and their adaptors, respectively (i.e. Grin2b, Grik1, Gria3, Grm5 and Grip1, Sharpin, Dlg2) (Figure S4K, highlighted pre-synaptic genes in pink and post-synaptic genes in green). Altogether these results suggest that considering the *Nova* family as a whole, *Nova2* (compared to *Nova1*) is the main driver of AS and importantly, may be most relevant for synaptic development of SST cINs.

### *SST-Nova1 and SST-Nova2* mutants have impaired afferent and efferent connectivity

To confirm our predictions from the AS analysis of conditional *Nova* mutants, we next sought to determine the effect of the loss of *Nova1* and *Nova2* on SST cIN synaptic development and function. To this end, we assessed the requirement for *Nova1* and/or *Nova2* for both the anatomical connectivity and physiological properties of SST cINs. *SST-dKO* mice were smaller in size and while generated at Mendelian ratios, many died as early as P8 and offspring often exhibited seizures (Figure S6A). In the single cKO mutants, we used IHC to quantify the density of SST cIN efferent synapses, defined as the apposition of VGAT+ (vesicle GABA transporter) and gephyrin+ puncta from an SST cIN axon within L1 of the S1 cortex at P8 (Figure 5A, black asterisks mark example puncta). We found that both *SST-Nova1* (0.281±0.041 puncta/um^2^ *SST-Nova1* vs 0.454±0.037 puncta/um^2^ ctl, pVal=0.003) and *SST-Nova2* (0.197±0.016 puncta/um^2^ *SST-Nova2* vs 0.454±0.037 puncta/um^2^ ctl, pVal=<0.0001) exhibited a significant reduction in SST+ synapses compared to control SST synapses within L1 (Figure 5B). To confirm the synaptic phenotype observed, we recorded the inhibitory outputs from SST cINs onto pyramidal cells in L2/3 and L5 using a conditional channelrhodopsin mouse line (*Ai32^F/F^*) crossed with *SST-Nova1*, *SST-Nova2, SST-Nova1/2-dKO* or SST-control mice (Figure 5C). We observed a significant reduction in the light evoked IPSC peak amplitude in *SST-Nova1* (283±36pA in *SST-Nova1* vs 623±120pA in ctl, pVal=0.0037, Figure 5D), *SST-Nova2* (340±85pA in *SST-Nova2 vs* 766±211pA ctl, pVal=0.0021, Figure 5E), and *SST-dKO* (248.9pA in *SST-dKO* vs 663pA in ctrl, Figure 5F) confirming that the anatomically observed reduction in synaptic output density is functionally significant in all mutants.

**Figure 4.**
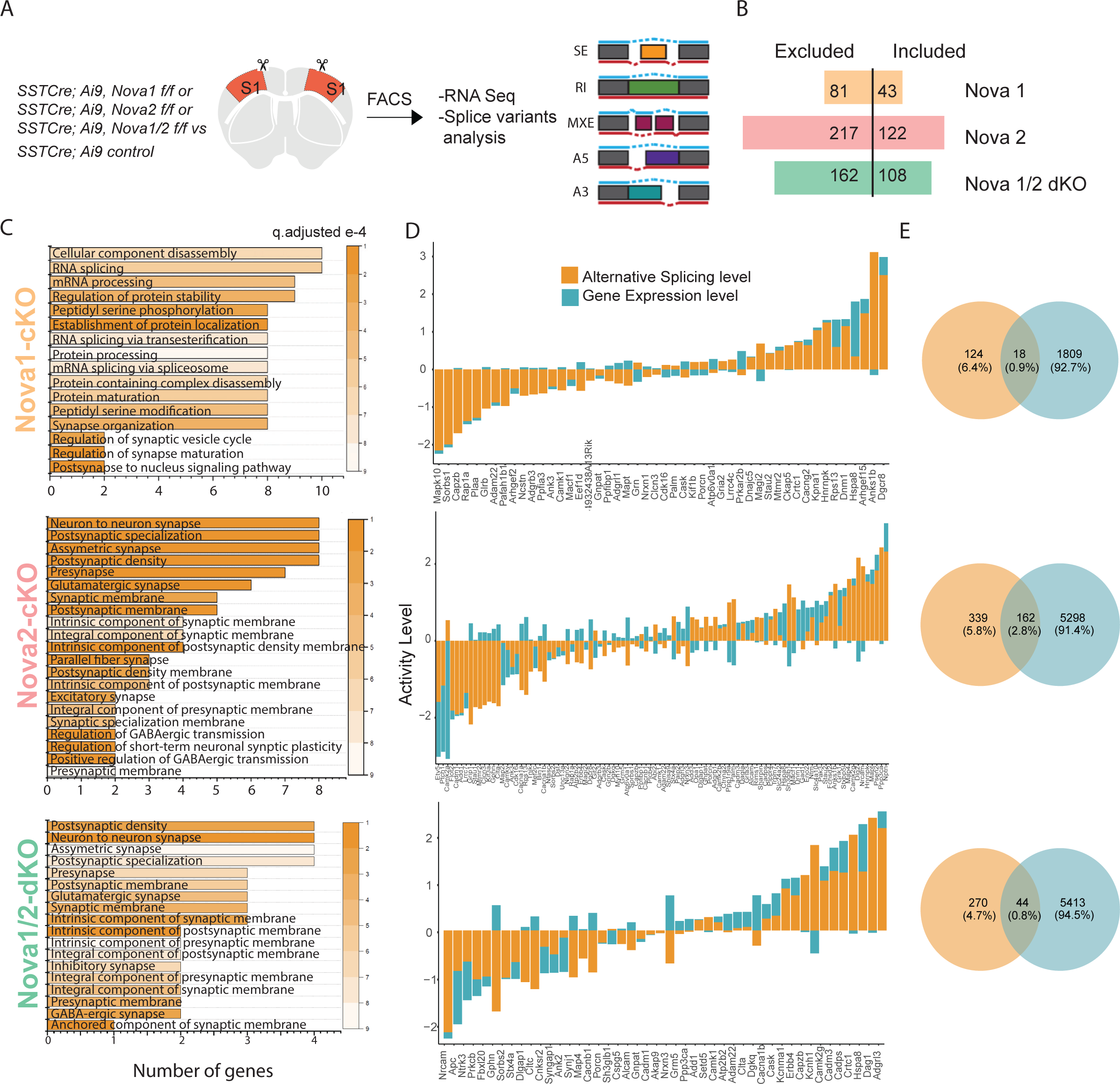

**Figure 5.**
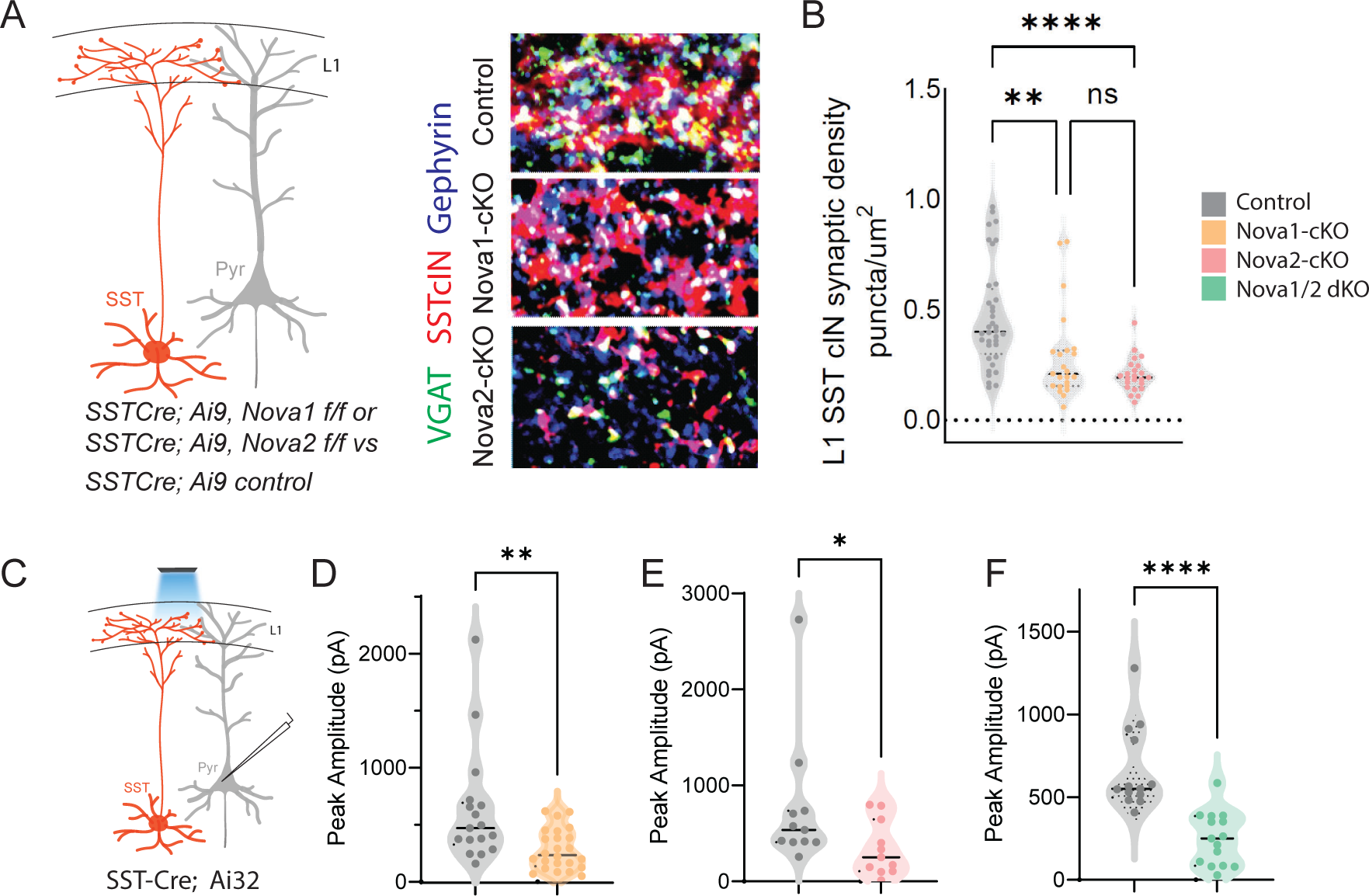
*SST-Nova1 and SST-Nova2* mutants have impaired afferent and efferent connectivity. A) SST+ cINs efferent structure: IHC of anti-RFP (red), anti-VGAT (green), and anti-Gephyrin (blue) to label the SST+ cIN axonal synaptic puncta (RFP+/VGAT+/Gephyrin+ puncta, white) in L1 S1 cortex of *SST-ctl, SST-Nova1,* and *SST-Nova2* mutant animals. B) Quantification of the density of SST+ cIN efferent synaptic puncta (RFP+/VGAT+/Gephyrin+) in L1 S1 cortex of *SST-ctl (n=26, S1 cortex from 3 mice), SST-Nova1 (n=26, S1 cortex, from 3 mice)* and *SST-Nova2 (n=15, S1 cortex from 3 mice)* mutant animals. **pVal=0.003, *SST-Nova1*; ***pVal<0.0001, *SST-Nova2* C) Schematic of channelrhodopsin (ChR2) experimental approach: SST-Cre control, SST-Nova1, SST-Nova2 or Nova1/2 (dKO) mutant mice were crossed with the Ai32 reporter line that expresses ChR2 in a Cre dependent manner. Blue light was delivered through the objective to record inhibitory response (IPSC) in neighboring excitatory neuron (grey). D-F) Quantification of the peak IPSC amplitudes recorded in excitatory neurons following SST stimulation in Nova1-cKO (D), Nova2-cKO (E) and Nova1/2-dKO (F) (n=20 cells from 3 mice each; **pVal=0.0037, *SST-Nova1*; **pVal=0.0021, *SST-Nova2,* ***pVal <0.001 ).

We also investigated whether the density of excitatory synapses onto SST cINs is affected by the loss of *Nova1* or *Nova2*. We performed IHC for Vglut1 (vesicular glutamate transporter) and Homer1c on *SST-Nova1* and *SST-Nova2* dendrites within the S1 cortex at P8 (Figure S6B, black asterisks mark example puncta). We quantified the density of putative excitatory synapses by the overlap of Vglut1+ and Homer1c+ puncta onto mCherry+ dendrites of SST cINs. We found that the number of putative excitatory afferent synapses onto *SST-Nova1* and *SST-Nova2* is significantly reduced compared to control SST cINs (0.144±0.016 puncta/um^2^ *SST-Nova1* vs 0.207±0.022 puncta/um^2^ ctl, pVal= 0.028 and 0.137±0.013 puncta/um^2^ *SST-Nova2* vs 0.207±0.022 puncta/um^2^ ctl, pVal= 0.012) (Figure S6C). To examine whether these anatomical abnormalities observed in *SST-Nova1* and *SST-Nova2* mutants affected synaptic function, we performed whole-cell patch clamp recordings to measure miniature excitatory postsynaptic currents (mEPSCs) within SST cINs (Figure S6D-I). In accordance with the puncta analysis, both *SST-Nova1* and *SST-Nova2* exhibited significant reductions in the mEPSC frequency (*SST-Nova1*: 1.16± 0.08 Hz vs *SST-Nova2*: 0.39±0.05 Hz vs ctl: 2.43±0.2 Hz, pVal=0.0025, Figure S6F). In addition, we observed a significantly increased mEPSC amplitude in *SST-Nova2* (*SST-Nova2*: -40±15.7pA vs *SST-Nova1*: -30.12±13.15pA vs ctl: -30.36±13.34pA, pVal=0.0.0001, Figure S6F right and S6H). Thus, while *SST-Nova2* cINs have a striking reduction in their excitatory inputs, the remaining excitatory synapses are functionally stronger than *Nova1* or control cINs. Moreover, the intrinsic properties of both *cKO* alleles were differentially affected. Specifically, we observed that the rheobase was significantly lower for *SST-Nova2* compared with either controls or *SST-Nova1* (*SST-Nova2*: 25±3pA vs ctl:120±25pA vs *SST-Nova1:* 70±15pA; pVal=0.01, Table S1). As rheobase is a measurement of the minimum current required to produce an action potential, *SST-Nova2* cIN*s* are potentially compensating for the loss of excitatory synapses by lowering the minimal current amplitude required for depolarization. Altogether these results solidify the role of both Nova1 and Nova2 in the synaptic development of SST cINs. Furthermore, consistent with the AS analysis, these results suggest that within SST cINs *Nova2* has a larger impact on the changes in synaptic connectivity compared to *Nova1*.

### Nova RNA binding proteins control activity dependent AS in SST cINS during development

Given that activity increases the expression level and nuclear localization of both Nova proteins, we hypothesized that their loss would result in changes in activity-dependent AS. To this end, we repeated our investigation of how Nova-dependent AS isoforms are altered in mutant mice. This time we examined the changes specifically following ECS within SST cINs during synaptogenesis *in vivo*. 2-3 hrs following ECS, we isolated SST cINs from *SST-dKO* mice (Figure 6A). Following augmentation of neuronal activity, we found that the loss of both *Nova* genes results in the differential splicing of 346 transcripts (FDR<0.05, |Δψ|≥0.1). These are comprised by 166 SE events (60 excluded and 106 included exons), 72 RI events (21 excluded and 51 included introns), 70 MXE events (33 excluded and 37 included exons), 9 A5 events (2 excluded and 7 included exons), and 29 A3 (20 excluded and 9 included exons) (Figure 6B). Many of these genes were categorized into synaptic gene ontology categories both in AS and GE data with a small degree of overlap (Figure 6C-E). As demonstrated previously, many synaptic genes exhibited higher AS change level compared to GE (Figure 6F and S7E). For example, the synaptic gene Nrxn1 was shown to have 4-fold difference in the AS level compared to GE (Figure 6F inset). Independent fluorescent RT-PCR amplifications with primers flanking the alternatively spliced segments confirmed the observed AS changes.

**Figure 6.**
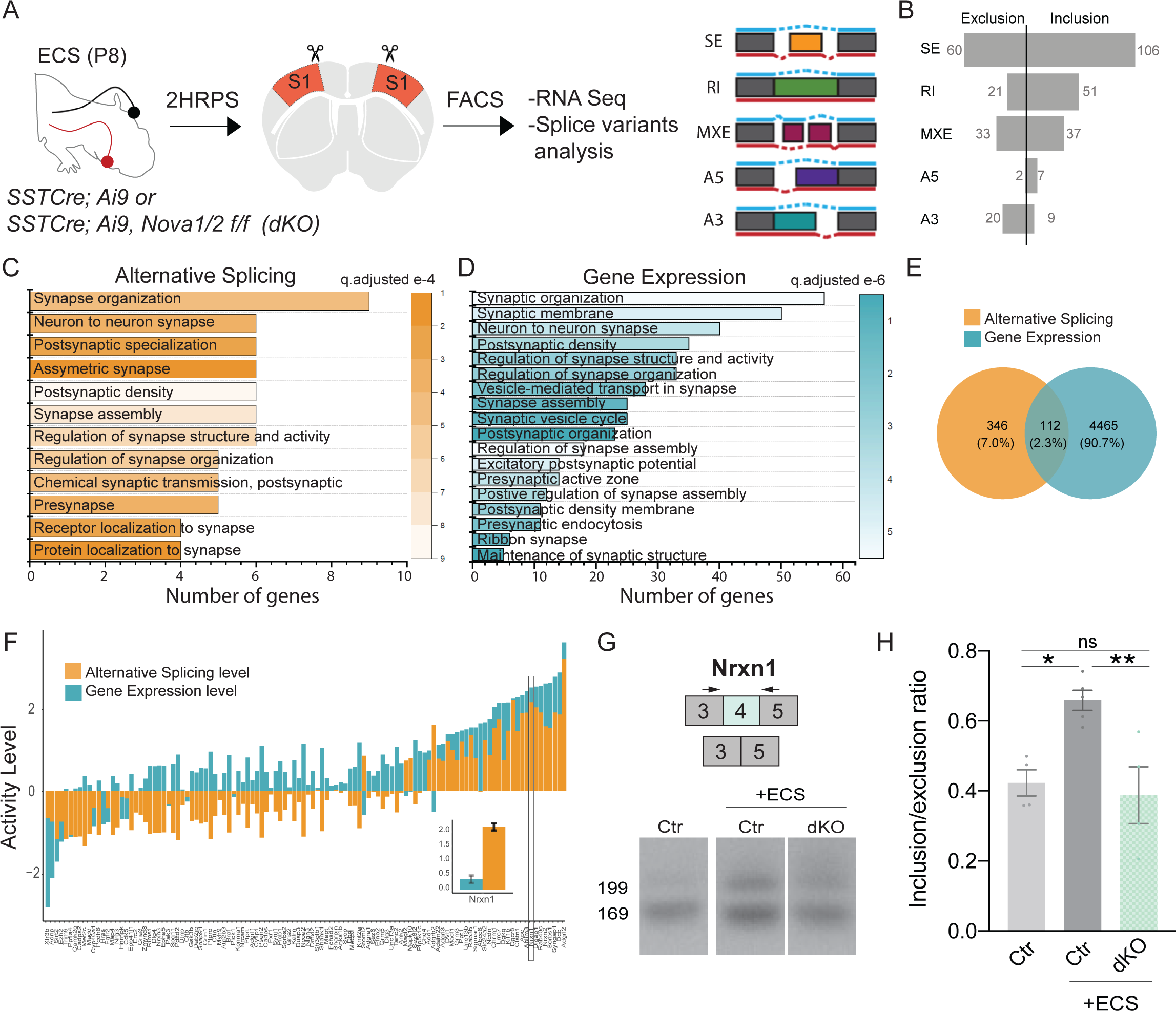
Nova RNA binding proteins control activity dependent AS in SST cINS during development. A) Schematic of experimental approach: Control and *SST-dKO* P8 animals were subjected to ECS then the S1 cortex was isolated to FACS purify SST+ cINs followed by RNAseq and splicing analysis. B) Magnitude of activity-dependent splicing changes within *SST-dKO* subjected to ECS compared to Ctr *SST-wt* cINs subjected to ECS (FDR <0.5, fold <0.1>), depicting 166 differential spliced SE (106 SE included, 60 SE excluded), 72 differential spliced RI (51 RI included, 21 RI excluded), 70 differential spliced MXE (37 included, 33 excluded), 9 differential spliced A5 (7 included, 2 excluded), 29 differential spliced A3 (9 included, 20 excluded). C) Synaptic gene ontology (GO) for the differentially spliced genes between ECS control vs ECS Nova1/2-dKO conditions. Color bar indicates adjusted q value. D) Synaptic gene ontology (GO) for the differentially expressed synaptic gene categories in the ECS control vs ECS Nova1/2-dKO conditions. E) Number and percentage of overlap between all differentially expressed genes (FC>0.5, pVal<0.05) and alternatively splice genes. F) Comparison of the activity level (Fold Change) of alternative splicing (AS) and gene expression (GE) amongst the shared genes that are both differentially expressed and differentially spliced. Inset shows that in the Nrxn1 gene AS level is larger (FC= 2.16) compared to the change in GE level (FC= 0.359). G) Example RT-PCR validation of alternative splicing (AS) events of activity- and Nova1/2-dependent alternative exon usage within the gene *Nrxn1* (top), Gel image of RT-PCR product from the amplification of exon 3 to exon 5 within *SST-ctl* cINs (Ctl) (left), ECS-treated Ctl (middle), and ECS-treated *SST-dKO* (right). H) Quantification of RT-PCR AS events of *Nrxn1.* *pVal=0.0194 Ctl vs Ctl+ECS; **pVal=0.0087 Ctl+ECS vs *SST-dKO*+ECS.

We were able to validate 70% of targets tested. For example, we validated the activity-dependent inclusion of exon 4 in *Nrxn1.* As predicted from RNAseq, SST cINs subjected to acute increases in activity from *SST-dKO* animals, compared to control SST cINs, exhibit a significant reduction in the expression of *Nrxn1* exon 4 (Figure 6G-H). Similarly, we validated the activity-dependent inclusion of exon 14 in *Syngap1,* a gene associated in multiple disorders including epilepsy and important for excitatory post-synaptic function (Figure S7C, D). Both activity-mediated gene expression and splicing changes are partially abolished by Nova1/2-dKO (Figure S7F-G). A list of exon coverage and inclusion levels for synaptic genes is presented in Table S2.

We found the majority of genes which undergo activity-induced Nova-dependent differential splicing were significantly enriched for GO categories such as pre-synaptic vesicular function, synapse organization, synaptic transmission, and neuronal growth (Figure 6C and S7A). Many of the genes within these categories are known to have important functions for axon organization and synaptogenesis such as, *Nrxn1, Nrxn3, PlxnA2,* and *EphA5*. Interestingly, the activity-dependent Nova AS targets were strikingly enriched for excitatory post-synaptic specializations such as, *Shank1*, *Syngap1, Dlg3, Grin1* and *Gria1*. Furthermore, these genes are predicted to function together in a direct PPI network representing specific pre-synaptic and post-synaptic biological processes (direct network pVal= 0.0009, 10000 permutations, Figure S7B). For example, the loss of *Nova* leads to an altered activity-dependent splicing program of multiple genes important to NMDA receptor-mediated signaling (Grin1) connected with PSD organization (i.e., Dlg3, Shank1) and Ca^2+^ -dependent signaling (i.e., Hras, Rapgef1) (Figure S7B).

In sum, the activity-mediated Nova-dependent AS changes within SST cINs are central for fine-tuning of synaptic development. We previously found that another important RNABP, Rbfox1, influences axonal development and also shuttles from the cytoplasm to the nucleus upon increase in activity in SST cINs (Lee et al., 2009; Wamsley et al., 2018). However, upon comparing the activity-dependent splicing programs within SST cINs of Rbfox1 (69 activity-dependent events) to Nova1/2 (346 activity-dependent events), we found Nova proteins control a much larger number of activity-dependent splicing events. This supports our hypothesis that Nova proteins are key players in the control of activity-dependent alternative splicing (Figure S8).

### Augmenting activity in Nova2 KO fails to enhance SST inhibitory output

Activity increases both the expression of Nova proteins as well as synapse formation, while conversely loss of Nova function causes a striking decrease in synaptogenesis and SST inhibitory output. Moreover, from our analysis of SST cIN cKOs, it was evident that of the two Nova proteins, *Nova2* has the more profound effect on the AS of genes involved in synaptogenesis. We therefore examined whether the loss of *Nova2* impaired the ability of augmented neuronal activity in SST cINs to promote the formation of efferent synaptic connectivity. To that end, we expressed NachBac in SST neurons in SST::Ai32 mice with *Nova2* deletions, compared with controls (Figure 7A). As previously shown, enhancing activity using NachBac resulted in increased Nova1/2 expression and localization into the nucleus in control mice (No Nova2-deletion, Figure 7A right). When we recorded from the pyramidal neurons in all conditions (control-No NachBac, control+NachBac, or N2-cKO+NachBac), we observed that enhancing activity in the Nova2-cKO did not result in an increase in inhibitory output of SST cINs (Figure 7B, right). This suggests that the activity dependent changes of synaptic strength depend upon the presence of Nova2.

**Figure 7.**
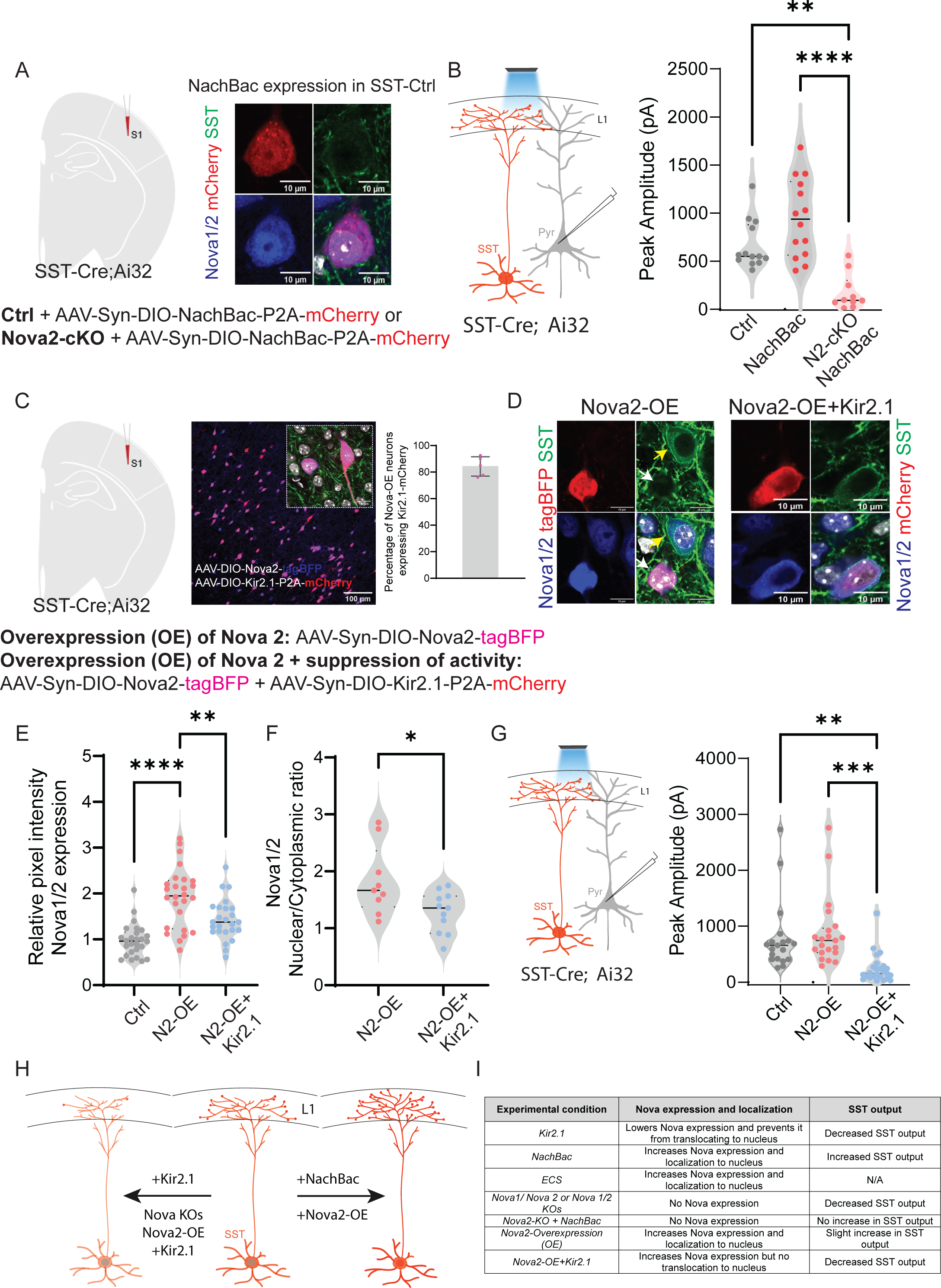
Augmenting activity in Nova2 KO fails to enhance SST inhibitory output. A) Left, experimental model: Injection of AAV-Syn-DIO-NachBac-P2A-mCherry (activating) in either control mice or SST::Nova2-cKO mice at P0 (analysis at P21). Right, example images showing the impact of NachBac activation (red) on Nova1/2 expression in controls. Note the translocation of Nova proteins (blue) to the nucleus (grey) in SST-cINs (green). Scale bar=10µm. B) Left, schematic of the recording scheme: SST-Cre::Ai32 optogenetic activation and recording from L5 pyramidal neurons. Right, quantification of the peak IPSC amplitude recorded from pyramidal neurons under no NachBac control conditions (grey), NachBac injections in SST-Ctrl animals (red dots) or NachBac injections in SST::Nova2-cKO animals (pink). (n=10-15 cells from each condition, N=3 mice; **≤ 0.01, ****≤ 0.001). C) Left, experimental model: Overexpression (OE) of Nova2 using the AAV-Syn-DIO-Nova2-tagBFP virus was injected into SST-Cre::Ai32 mice either alone or while suppressing activity using Kir2.1 (AAV-Syn-DIO-Kir2.1-P2A-mCherry. Middle, an image showing the co-expression of Nova2-tagBFP (blue) and Kir2.1-mCherry (red). Inset shows co-localization of both proteins in SST neurons. Scale bar of inset=10um. Right, percentage of overlap between the two viruses in SST neurons, quantified as percentage of Nova2-OE neurons that also express Kir2.1-mCherry (∼85%). D) Left panels, Representative images of IHC against tagBFP (red), and Nova1/2 (anti-Nova1/2, blue) in SST-Nova2OE cells in SST-Cre::Ai32 mice (green labels Ai32 expression). Right panels, SST-Nova2OE+KIR2.1 cell. Bottom right panels represent merged images. Note the exclusion of Nova proteins from the nucleus in Nova2-OE+Kir2.1 conditions. E) Quantification of the relative pixel intensity of Nova1/2 expression in SST cINs (n=25/26 cells for each condition, pVal=**≤ 0.01, ****≤ 0.001). F) Quantification of the Ratio of Nova1/2 localization within the nucleus to cytoplasm from Nova2OE SSt cINs (pink) and Nova2OE+KIR2.1 (blue). (n=10 cells from 3 mice; pVal=*≤ 0.05) G) Right, recording schematic. Left, Peak IPSC amplitude recorded from pyramidal neurons in response to optogenetic stimulation of SST-cINs in either the Nova2-OE condition or Nova2-OE+Kir2.1 condition (n=19-25 cells in each condition, pVal=**≤ 0.01, ***≤ 0.005). H) Model of experimental findings: center is a cartoon wildtype SST cIN depicting normal expression of Nova1/2 with the soma (red) whereas, on the left, the conditional loss of *Nova1, Nova2,* or the expression of KIR2.1 alone or dual overexpression of Nova2 and KIR2.1 results in the reduction in Nova expression and restricts Nova localization to the cytoplasm (In the case of cKO animals the protein is lost completely). This effect is accompanied by a reduction in the connectivity of SST cINs. To the contrary, Expression of NaChBac and/or overexpression of Nova2 alone results in expression of Nova throughout the cell and nucleus and is accompanied by an increase in the SST cINs output. I) Summary table of experimental findings in all conditions tested.

Conversely, we examined whether over-expression (OE) of *Nova2* alone could phenocopy the observed changes in connectivity within SST cINs and whether that was affected by reducing the activity level of the cell (using Kir2.1). To that end, we either overexpressed *Nova2* alone specifically in SST+ neurons using an AAV virus (AAV-Syn-Nova2-P2A-mCherry) in *SST^Cre^:* mice within the S1 cortex or in conjunction with Kir2.1 OE (Figure 7C). As in the case of increasing activity (either constitutively, NaChBac, or acutely, ECS), the nuclear localization of Nova was robustly increased when *Nova2* was overexpressed in the SST cINs (Figure 7D-F). The increased nuclear localization of Nova that was observed with the *Nova2* OE was abolished when the activity of the cells was cell-autonomously reduced using KIR2.1.

We next also examined whether suppressing activity while overexpressing *Nova2* (Nova2-OE) impacts the inhibitory output of SST neurons (Figure 7G left). The dual expression of Nova2 OE and KIR2.1 within SST cINs prevented the small increase of peak IPSC amplitude observed with Nova2-OE alone. Perhaps most strikingly, as with our initial KIR2.1 experiment, the levels of Nova2 protein despite being constitutively OE were reduced in cells co-expressing KIR2.1 (Figure SE). This provides strong evidence that the stability and nuclear localization of Nova protein is dependent on the level of basal activity within SST cINs. Therefore, a certain level of activity is needed to maintain Nova protein function, and conversely, Nova proteins are needed to mediate activity dependent changes in alternative splicing of synaptic proteins.

## Discussion

In the present study, we have examined the interacting contributions of neuronal activity and the Nova RNABPs on synaptogenesis of SST cINs. Our analysis began with the observation that activity levels strongly influence the maturation of SST cINs. Acutely evoking activity during circuit integration with ECS resulted in both transcriptional and translational upregulation of Nova proteins and promoted their localization to the nucleus. This was accompanied by a striking change in both the GE and AS of synaptic genes and culminated in enhanced synaptogenesis within SST cINs. We then systematically examined the interdependence between these three observations.

Our results indicate that during circuit formation, activity levels within SST cINs correlate with changes in AS and together act to regulate the formation of afferent/efferent connectivity. These events appear to be tightly linked to Nova function, as the expression, localization and splicing activity of both Nova1 and Nova2 proteins are strongly modulated by activity. Examination of how splicing events are impacted by *Nova* single and compound cKOs in SST cINs demonstrates that developmental RNA splicing events in these cells are particularly impacted by the loss of *Nova2.* This is mirrored by the magnitude in reduction of excitatory input and inhibitory output within *Nova2* null SST cINs, as evidenced by a structural and functional decrease in their synaptic contacts. The relationship between activity and Nova2 function during development is interdependent. During these periods, boosting activity cell autonomously within Nova2-cKOs fails to increase the structural or physiological output of SST-cINs. Conversely, over-expression of *Nova2* in SST cINs can enhance these activities but this phenomenon can be suppressed by simultaneous dampening their excitability. Together these findings demonstrate that activity is coupled to synaptogenesis in SST cINs by a mechanism involving Nova proteins. Whether these effects are regulated through their contributions to AS, GE or a combination of both remains to be determined.

With regards to AS in particular, Nova function is a core regulator of alternative splicing in many cell types, including SST cINs. It however represents only one of a host of RNABPs within the CNS. Indeed, a recent study demonstrated that within the mature brain many classes of neurons, including SST cINs, can be classified both by their expression levels of RNABPs and their corresponding repertoire of alternatively spliced mRNAs (Furlanis et al., 2019). Comparison of this work to our present findings illustrate that both the expression of RNABPs and the patterns of AS are strongly regulated across development, a phenomenon that may reflect developmental changes in neuronal activity. Consistent with this RNA binding splice factors have previously been shown to promote alternative splicing of synaptic proteins in response to neuronal depolarization and Ca^2+^ signaling (Eom et al., 2013; Mauger et al., 2016; Quesnel-Vallières et al., 2016; Vuong et al., 2016), For example, previous research demonstrated that the splicing of neurexins, a gene family known to function in synaptogenesis, are mediated through the actions of the SAM68 splicing factor (Iijima et al., 2011b). Similarly, It has also been illustrated that neuronal activity reduces the expression of the SRRM4 RNA-binding protein, which resulted in altered RNA splicing and a corresponding decrease in excitatory synapses (Quesnel-Vallières et al., 2016). As such AS represents a largely unexplored but central genetic mechanism, capable of directing cell-type development and synaptic formation specifically.

Understanding both the repertoire of splice factors and the cell-specific patterns of splicing across development will undoubtedly provide further insight into how AS influences cIN development. One could imagine systematically examining the role of these differential splice mRNA variants through combinatorial knockdown or over-expression. However, this would face enormous technical challenges, even if restricted to only those that are Nova-dependent. As we show here many of these genes have been shown to function together (PPI networks). As such AS appears to coordinately target specific biological mechanisms. Given that the abundance of the specific splice forms of different genes within SST cINs is relative rather than absolute, it appears that AS has been coopted by development as an effective mechanism to fine-tune particular biological phenomena. The flexibility of AS to regulate the composition and levels of particular genes allows cells to adjust their biological function in accordance with both their identity and state (i.e. developmental period, neuronal activity, etc). As a result, the abundance of particular splice forms co-varies as a function of both transcription and AS. Taken together, this argues that conditional removal of RNABPs, such as Nova2, provides an effective approach for understanding the role of AS within discrete cell types. Additionally, Nova proteins have a yet unexplored role in regulating gene expression, most likely through their ability to regulate the stability of RNA molecules.

In sum, our results show a clear interdependence between activity, Nova function and synaptic formation/strength in SST cINs. The interaction between activity and Nova function is bidirectional. Activity regulates the RNA, protein levels and intracellular localization of Nova proteins within SST cINs, while Nova proteins are in turn required for the activity-dependent regulation of synaptic formation and function (see model Figure 7H). When SST cIN activity is increased with ECS or with NaChBac expression, *Nova* transcripts as well as protein are upregulated and shuttled to the nucleus. The mechanisms for activity-dependent changes in Nova expression and localization are unknown. It is possible that the *Nova* gene loci may contain binding sites for immediate-early-genes (i.e. cFOS, Jak/Jun, EGF) or specific activity-dependent transcription factors (i.e. NPAS4, Satb1). With regard to control of its localization, previous work has discovered a nuclear-localization signal (NLS) within the Nova protein domains. It is however unknown whether their activation is also mobilized by splicing or post-translational modifications. For instance, *Rbfox1* undergoes activity-dependent mRNA splicing that results in exposure of an NLS and localization to the nucleus (Lee et al., 2009; Wamsley et al., 2018). Furthermore, our results indicate that activity itself regulates Nova2 RNA and protein stability. In the presence of KIR2.1, the levels of Nova protein appear to be dramatically reduced, even when *Nova2* is over-expressed. In this latter context, clearly *Nova2* levels are not constrained by mRNA production. These results indicate that the stability of Nova protein is at least partly dependent on activity. Taken together, these findings indicate that there exist multiple mechanisms by which cell activity is coupled to Nova function and AS within SST cINs.

We and others have shown that activity regulates programmed cell death (Priya et al. 2018; Denaxa et al. 2018; Wong et al. 2018). However, we observed no indication that the loss of *Nova2* impacted SST cIN survival. In addition, we observed that NaChBac and KIR2.1 could modulate synaptogenesis in SST cINs both during and after the peak of cell death in this region (data not shown). Conversely, the number of phenotypic changes observed in conditional *Nova* loss of function mutants suggests that these genes have effects beyond synaptogenesis. Nova2, in particular, also targets genes involved in protein trafficking to the membrane, cell-cell signaling, and neurotransmitter/ion channel function, indicating it influences multiple aspects of SST cIN maturation. In addition, prior work from the Darnell lab has demonstrated a role for Nova2 in both migration and axonal pathfinding within the cortex, spinal cord and brain stem (Saito et al., 2016; Yano et al., 2010). Taken together clearly much remains to be understood concerning the role Nova proteins play during development in specific brain regions, circuits, and cell types. Indeed, given the broad expression of Nova proteins and the strong phenotypes associated with both conditional and global *Nova* loss of function, studies of this RNABP will no doubt provide further insights into their contribution to normal and disease brain function.

## CONTACT FOR REAGENT AND RESOURCE SHARING

Please contact GF for reagents and resources generated in this study.

## EXPERIMENTAL MODEL AND SUBJECT DETAILS

### • Mouse maintenance and mouse strains

All experimental procedures were conducted in accordance with the National Institutes of Health guidelines and were approved by the Institutional Animal Care and Use Committee of the NYU School of Medicine and Harvard Medical School. Generation and genotyping of, *SST^Cre^* (Taniguchi et al., 2011), *RCE^eGFP^*(Sousa et al., 2009), *Lhx6* BAC transgenic (referred to as tg*Lhx6;eGFP*) (Gong et al., 2003), *Nova1^LoxP/LoxP^* (Yuan et al., 2018a), *Nova2^LoxP/Lox^*(Saito et al., 2019), TRE-Bi-SypGFP-TdTomato (Li et al., 2010), and Ai9 *^LoxP/LoxP^* Ai32 *^LoxP/LoxP^*, *Rosa-tTa ^LoxP/LoxP^* (commercially available from Jax laborites). All mouse strains were maintained on a mixed background (Swiss Webster and C57/ B16). The day of birth is considered P0. Information about the mouse strains including genotyping protocols can be found at http://www.jax.org/ and elsewhere (see above references).

## METHOD DETAILS

### • Immunochemistry and imaging

Embryos, neonate, juvenile and adult mice were perfused inter cardiac with ice cold 4% PFA after being anesthetized on ice (neonates) or using Sleepaway injection. Brains that were processed for immunofluorescence on slides were post-fixed and cryopreserved in 30% sucrose. 16µm coronal sections were obtained using Cryostat (Leica Biosystems) and collected on super-frost coated slides, then allowed to dry and stored at -20°C until use. For immunofluorescence, cryosections were thawed and allowed to dry for 5-10 min and rinsed in 1x PBS. They were incubated at room temperature in a blocking solution of PBST (PBS-0.1%Tx-100) and 10% normal donkey serum (NDS) for 1hr, followed by incubation with primary antibodies in PBS-T and 1% NDS at 4°C overnight or 2 days. Samples were then washed 4 times with PBS-T and incubated with fluorescence conjugated secondary Alexa antibodies (Life Technologies) in PBS-T with 1% NDS at room temperature for 1 hr. Slides were incubated for 5min with DAPI, washed 3 times with PBS-T. Then slides were mounted with Fluoromount G (Southern Biotech) and imaged.

Brains that were processed for free-floating immunofluorescence were first post-fixed in 4% PFA overnight at 4°C. 50 µm-thickness brain slices were taken on a Leica vibratome and stored in a cryoprotecting solution (40% PBS, 30% glycerol and 30% ethylene glycol) at -20°C. For immunofluorescence, floating sections were blocked for 1hr at RT in normal donkey or goat serum blocking buffer and incubated for 2-3 days at 4°C with primary antibodies in blocking buffer. Sections were washed 4 x 30 min at RT in PBST, incubated overnight at 4°C with secondary antibodies and DAPI in blocking buffer, washed 4 x 30 min at RT in PBST before being mounted on super-frost plus glass slides. Primary antibodies are listed in Key Resource Table.

### • Nova1/2 Localization

To quantify the Nova localization in SST cINs, mCherry+/SST cIN, KIR2.1+/SST cINs or NaChBac+/SST cIN (n=27 cells from 3 mice each); control/SST cIN or ECS+/SST cINs (n=27 cells from 3 mice each); Nova2OE/SST cIN or Nova2OE+KIR2.1/SST cINs (n=20 cells from 3 mice) were binned into two categories based on the cell compartment Nova1/2 protein was localized to: Cytoplasmic restricted or Nuclear-expressing (comprised of nuclear restricted or whole soma localization). The number of Nuclear-expressing cells was then divided by the number of cytoplasmic restricted cells to obtain a ratio for Nova localization from either mCherry+/SST cIN or KIR2.1+/SST cINs. This was collected from at least three tissue sections from at least 3 animals.

### • Electroconvulsive Shock

Electroconvulsive stimulation (ECS) was administered to animals with pulses consisting of 1.0 s, 50 Hz, 75 mA stimulus of 0.7 ms delivered using the Ugo Basile ECT unit (Model 57800, as previously described (Guo et al., 2011; Ma et al., 2009). Control/sham animals were similarly handled using the exact same procedure but without the current administration.

### • Confocal imaging and synaptic puncta analysis

Animals were perfused as described above. Post-fixation incubation prior to cryopreservation was skipped. Cryostat sections (16 μm) were subjected to IHC as described above. Images were taken within the S1 cortex of at least three different sections from at least three different animals per genotype with a Zeiss LSM 800 laser scanning confocal microscope. Scans were performed to obtain 4 optical Z-sections of 0.33 μm each (totaling ∼1.2μm max projection) with a 63x/1.4 Oil DIC objective. The same scanning parameters (i.e. pinhole diameter, laser power/offset, speed/averaging) were used for all images. Maximum projections of 4 consecutive 0.33μm stacks were analyzed with ImageJ (NIH) puncta analyzer plugin (Ippolito and Eroglu, 2010) to count the number of individual puncta consisting of pre-synaptic and post-synaptic markers that are close enough together to be considered a putative synaptic puncta. Synaptic puncta density per image was calculated by normalization to total puncta acquired for each individual channel accounted in each image for each condition. Puncta Analyzer plugin is written by Barry Wark, and is available for download (https://github.com/physion/puncta-analyzer).

Nova protein intensity was performed as: Cryostat sections of 20 um were immunostained with goat anti-mCherry and human anti-pan Nova (from Darnell Lab). Images were analyzed using Fiji/ImageJ and Nova1/2 protein intensity levels were assessed normalized against area of the cells expressing the AAV.

### • Electrophysiological recordings

Slice preparation: Acute brain slices (300 μm thick) were prepared from P18-P22 mice. Mice were deeply anesthetized with isofluorane. The brain was removed and placed in ice-cold modified artificial cerebrospinal fluid (ACSF) of the following composition (in mM): 87 NaCl, 26 NaHCO3, 2.5 KCl, 1.25 NaH2PO4, 0.5 CaCl, 4 MgCl2, 10 glucose, 75 sucrose saturated with 95% O2, 5% CO2 at pH=7.4. Coronal sections were cut using a vibratome (Leica, VT 1200S). Slices were then incubated at 34C for 30 minutes and then stored at room temperature until use.

Recordings: Slices were transferred to the recording chamber of an up-right microscope (Zeiss Axioskop) equipped with IR DIC. Cells were visualized using a 40X IR water immersion objective. Slices were perfused with ACSF of the following composition (in mM): 125 NaCl, 25 NaHCO3, 2.5 KCl, 1.25 NaH2PO4, 2 CaCl2, 1 MgCl2, 20 glucose, saturated with 95% O2, 5% CO2 at pH=7.4 and maintained at a constant temperature (31°C) using a heating chamber. Whole-cell recordings were made from randomly selected tdTomato-positive SST interneurons or tdTomato negative pyramidal cells from layer II-III or layer V of the somatosensory cortex. Recording pipettes were pulled from borosilicate glass capillaries (Harvard Apparatus) and had a resistance of 3-5 MΩ when filled with the appropriate internal solution, as reported below. Recordings were performed using a Multiclamp 700B amplifier (Molecular Devices). The current clamp signals were filtered at 10 KHz and digitized at 40 kHz using a Digidata 1550A and the Clampex 10 program suite (Molecular Devices). Miniature synaptic currents were filtered at 3 kHz and recorded with a sampling rate of 10 kHz. Voltage-clamp recordings were performed at a holding potential of 0mV. Current-clamp recordings were performed at a holding potential of -70 mV. Cells were only accepted for analysis if the initial series resistance was less than 40 MΩ and did not change by more than 20% throughout the recording period. The series resistance was compensated online by at least ∼60% in voltage-clamp mode. No correction was made for the junction potential between the pipette and the ACSF.

Passive and active membrane properties were recorded in current clamp mode by applying a series of hyperpolarizing and depolarizing current steps and the analysis was done in Clampfit (Molecular Devices). The cell input resistance was calculated from the peak of the voltage response to a 50 pA hyperpolarizing 1 sec long current step according to Ohm’s law. Analysis of the action potential properties was done on the first spike observed during a series of depolarizing steps. Threshold was defined as the voltage at the point when the slope first exceeds a value of 20 V.s-1. Rheobase was defined as the amplitude of the first depolarizing current step at which firing was observed. Analysis of miniature inhibitory events was done using Clampfit’s template search.

Pipette solutions: Solution for voltage-clamp recordings from pyramidal cells (in mM): 125 Cs-gluconate, 2 CsCl, 10 HEPES, 1 EGTA, 4 MgATP, 0.3 Na-GTP, 8 Phosphocreatine-Tris, 1 QX-314-Cl and 0.4% biocytin, equilibrated with CsOH at pH=7.3. Solution for current clamp recordings from SST cINs (in mM): 130 K-Gluconate, 10 KCl, 10 HEPES, EGTA, 4 MgATP, 0.3 NaGTP, 5 Phosphocreatine and 0.4% biocytin, equilibrated with KOH CO2 to a pH=7.3.

### • *Nova2* OE/ *Nova2* OE +KIR2.1 Experiment

*SST^Cre^* mice crossed with *Ai32* mice were injected at P0/1 with either AAV2/1-Syn-DIO-Nova2-tagBFP or together with AAV2/1-Syn-DIO-Kir2.1-mCherry at 1:1 ratio in the S1 cortex. Mice were perfused at P21, brains harvested, sucrose protected and sectioned on a freezing microtome (Leica) at 20um thickness as described above. Primary antibodies are listed in Key Resource Table.

### • Optogenetic stimulation

Blue-light (470 nm) was transmitted to the slice from an LED placed under the condenser of an up-right microscope (Olympus BX50). IPSCs were elicited by applying single 1 ms blue-light pulses of varying intensities (max. stimulation intensity ∼0.33 mW/mm^2^) and directed to L2/3 or L5 of the slice in the recording chamber. Light pulses were delivered every 5 seconds. The LED output was driven by a TTL output from the Clampex software of the pCLAMP 9.0 program suite (Molecular Devices).

### • Isolation of cortical interneurons from the developing mouse cerebral cortex

Cortical interneurons were dissociated from postnatal mouse cortices (P8) as described (Wamsley et al., 2018). We collected at least 3-5 cKO and 3-5 *ctl* brains and maintained overall balanced numbers of females and males within each condition, in order to avoid sex-related gene expression biases. Following dissociation, cortical neurons in suspension were filtered and GFP+ or TdTomato+ fate-mapped interneurons were sorted by fluorescence activated-cell sorting (FACS) on either a Beckman Coulter MoFlo (Cytomation), BD FACSAria II SORP or Sony SY3200. Sorted cINs were collected and lyzed in 200µl TRIzol LS Reagent, then thoroughly mixed and stored at -80°C until further total RNA extraction.

### • Nucleic acid extraction, RNA amplification, cDNA library preparation and RNA sequencing

Total RNAs from sorted SST cINs (P8 mouse S1 cortices for Figure 2, Figure S2, Figure S4, Figure S6 and Figures 5) were extracted using TRIzol LS Reagent and PicoPure columns (if <20K cells were recovered) or PureLink RNA Mini Kit (if >20K cells were recovered), with PureLink DNase for on-column treatment, following the manufacturers’ guidelines. RNA quality and quantity were measured with a Picochip using an Agilent Bioanalyzer and only samples with high quality total RNA were used (RIN: 7-10). 20ng of total RNA was used for cDNA synthesis and amplification, using NuGEN Ovation RNA-Seq System V2 kit (NuGEN part # 7102). 100 ng of amplified cDNA were used to make a library using the Ovation Ultralow Library System (NuGEN part # 0330). The samples were mulitplexed and subjected to 50-nucleotide paired-end read rapid with the Illumina HiSeq 2500 sequencer (v4 chemistry), to generate >50 million reads per sample. Library preparation, quantification, pooling, clustering and sequencing was carried out at the NYULMC Genome Technology Center. qRT-PCR (quantitative RT-PCR) was performed using SYBR select master mix (Thermo-Fisher Scientific) on cDNA synthesized using SuperScript II reverse transcriptase and oligo(dT) primers.

List of RT- and qRT-PCR primers:

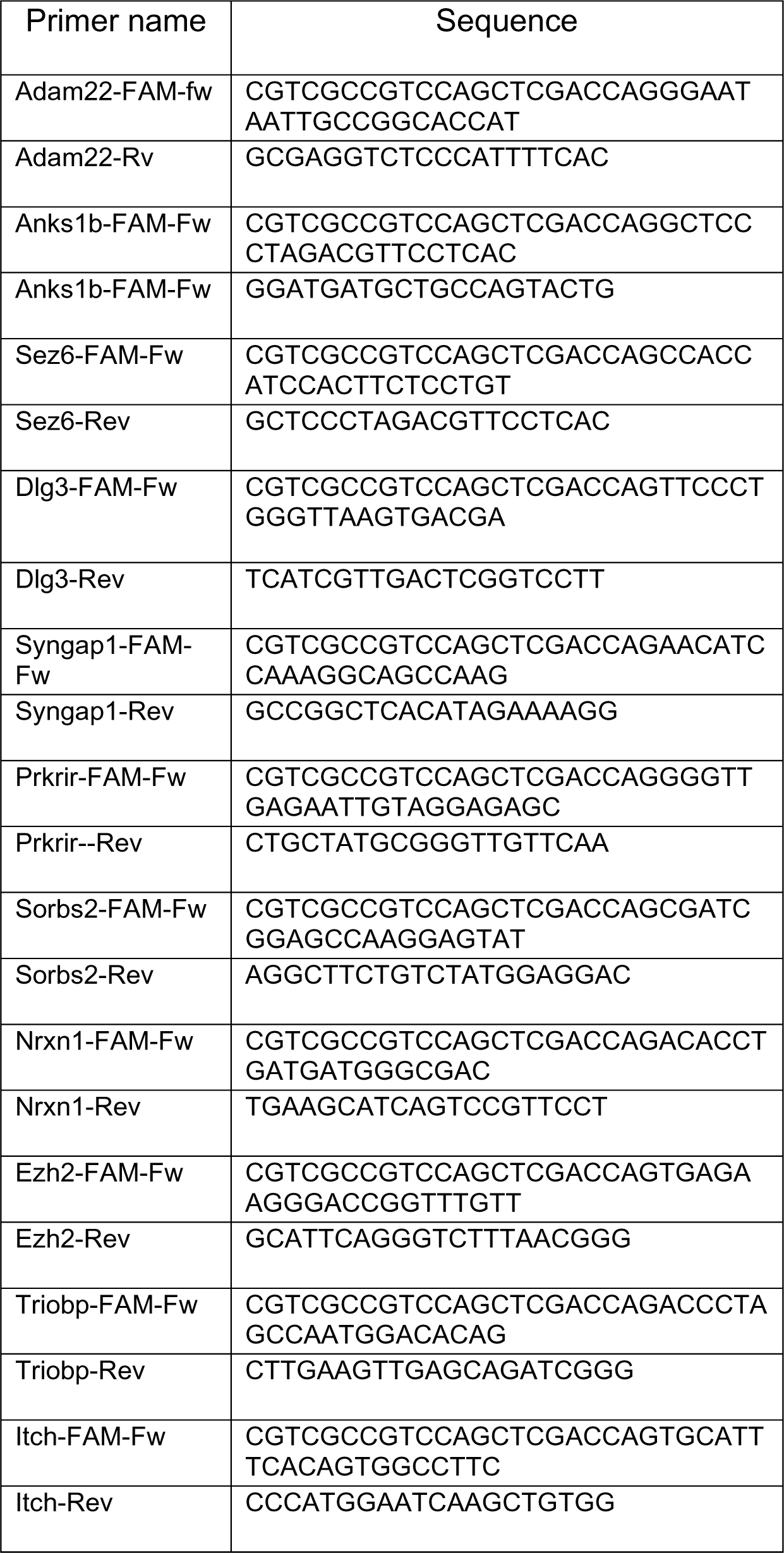

### • Bioinformatics

Downstream computational analysis were performed at the NYULMC Genome Technology Center and at KAUST. All the reads were mapped to the mouse reference genome (mm10) using the STAR aligner (Dobin et al., 2013). Quality control of RNAseq libraries (i.e. the mean read insert sizes and their standard deviations) was calculated using Picard tools (v.1.126) (http://broadinstitute.github.io/picard/). The Read Per Million (RPM) normalized BigWig files were generated using BEDTools (v2.17.0) (Quinlan and Hall, 2010) and bedGraphToBigWig tool (v4). For the SST cIN P8 ECS, approx. 60E6-80E6 reads were aligned per sample; for P8 SST-Nova1, SST-Nova2, SST-dKO, approx. 60E6-70E6 reads were aligned per sample; for P8 SST-cIN wt ECS and SST-Nova-dKO ECS, approx. 60E6-80E6 reads were aligned per sample. The samples processed for downstream analysis were as follows: 9 samples for SST cIN +ECS versus SST cIN ctl at P8 (4/5 samples per condition), 6 samples for *SST-Nova1* removal versus SST cIN ctl (3 samples per genotype), 6 samples for *SST-Nova2* removal versus SST cIN ctl (3 samples per genotype), 6 samples for *SST-dKO* removal versus SST cIN ctl (3 samples per genotype) and 7 samples for *SST-dKO* removal ECS versus SST cIN ctl ECS (4 control samples, 3 cKO samples). We performed differential expression analysis using DESeq2 R package for calculating the ecpression level of transcripts between different conditions. Genes with an adjusted p value <0.05 and log fold change (FC)≥ 0.5 were considered differentially expressed.

We used rMATS (v3.0.9) (Shen et al., 2014) to quantify the AS event types (i.e.skipped exons (SE), alternative 3’ splice sites (A3SS), alternative 5’ splice sites (A5SS), mutually exclusive exons (MXE) and retained introns (RI)). rMATS uses a counts-based model, it detects AS events using splice junction and exon body counts and calculates an exon inclusion level value ψ for each event in each condition. It then determines the differential |Δψ| value across conditions (cut-offs for significance were placed at *FDR*<0.05 and |Δψ|≥0.1). To compare the level of similarity among the samples and their replicates, we used two methods: classical multidimensional scaling or principal-component analysis and Euclidean distance-based sample clustering. The downstream statistical analyses and generating plots were performed in Rstudio (Version 1.1.456) (http://www.r-project.org/).

To assess the enrichment for the Nova-binding motif in the differentially regulated exons we utilized rMAPS (Park et al., 2016). We utilized the raw output from rMATS analysis (6 RNAseq experiments of SST cINs +ECS vs SST cINs ctl) with significant splicing events cut off at FDR>50%. rMAPS performs position weight analysis to assess the enrichment of RNA binding protein binding motifs in the exonic and flanking intronic regions of up-regulated or down-regulated exons and plots the motif density along with a given pValue in comparison to unregulated exons.

We performed GO analysis using the DAVID online Bioinformatics Resources 6.8 at FDR >0.05 (unless otherwise specified) (Huang et al., 2008) and tested PPI networks by utilizing DAPPLE at 10,000 permutations (Rossin et al., 2011). The GO categories were assigned to each group of genes, and after that, we used ClusterProfiler, the R function that helps with gene functional annotation and to perform GO enrichment analysis.

### • Validation of SST-cINs AS activity-dependent exons by RT-PCR

Total RNAs from sorted cINs from wt/ctl SST cINs, ECS SST cINs, and ECS SST-dKO were extracted as described above and at least three independent biological replicates were used in each experiment. RT-PCR validation of regulated exons was performed as described before (Han et al., 2014). After denaturation, samples were run on 10% Novex™ TBE-Urea Gels (ThermoFisher). Gels were directly scanned by ChemiDoc™ Imaging System (Bio-Rad) and quantified by ImageStudio program (Licor).

## QUANTIFICATION AND STATISTICAL ANALYSIS

No statistical method was used to pre-determine sample sizes, but our sample sizes were similar to those reported in previous publications in the field. In all figures: *, *p-*value<0.05; **, *p-*value<0.01; ***, *p-*value<0.001; ****, *p-*value<0.0001. Statistical analyses for motif enrichment was performed by rMAPS and differential alternative splicing changes were performed using rMATS. Percentages were compared with repeated t-tests in *GraphPad Prism* or *Rstudio*, and means ± (standard deviation, SD) are represented. Some statistical analyses and generating plots were performed in R environment (v3.1.1) (http://www.r-project.org/).

All values presented in the manuscript are average ± standard error of the mean (SEM). The statistical values for the intrinsic physiology are obtained using one-way ANOVA with Bonferroni correction for multiple comparisons between the different genotypes: Controls, Nova1-cKO, Nova2-cKO and Nova-dKO (*p ≤ 0.05, **p ≤ 0.01, **p ≤ 0.005). For the Channelrhodopsin output, we first determined if the data is normally distributed using Lilliefors test. In case of normal distribution, we performed student’s t-test was used to compare Control vs Nova1-cKO, and Control vs Nova2-cKO (*p ≤ 0.05, **p ≤ 0.01, ***p ≤ 0.005).

## DATA AND SOFTWARE AVAILABILITY

The accession number for the RNA sequencing data reported in this paper is NCBI GEO GSE143316. All processed RNA sequencing and splicing analysis data and sashimi plots can also be found at https://github.com/IbrahimLab-23/Nova-proteins-and-synaptic-integration-of-Sst-interneurons

## Acknowledgements

We would like to thank the NYULMC Division of Advanced Research Technologies and their personnel: Mouse Genotyping Core (Jiali Deng and Jisen Dai); Cytometry and Cell Sorting Core (Kamilah Ryan, Keith Kobylarz, Yulia Chupalova, and Michael Gregory); Genome Technology Center (Adriana Heguy); and Applied Bioinformatics Laboratory (Aristotelis Tsirigos), which is supported in part by grant UL1 TR00038 from the National Center for Advancing Translational Sciences (NCATS), NIH. CCSC and GTC are supported by the Cancer Center Support Grant, <u>P30CA016087</u>, at the Laura and Isaac Perlmutter Cancer Center. We would also like to thank Yanjie Qiu and Marian Fernandez-Otero for helping with genotyping at Harvard Medical School. We would like to thank the extended Fishell Laboratory for critical reading of the manuscript. Work in the G.F. lab is supported by the following NIH grants: R01 NS081297, R01 MH071679, UG3 MH120096, P01 NS074972 and by the Simons Foundation SFARI.

## Author Contributions

L.A.I, B.W, and G.F. conceived and developed the methodology and project. B.W., L.A. and N.Y. performed histology, behavior, imaging, and image analysis experiments. N.Y. performed biochemistry experiments with assistance from E.F. N.A.G, A.K.-J. and B.W. analyzed the RNA-seq experiments. L.A.I. and E.S performed electrophysiological experiments. L.A. and N.Y. performed the Activity-dependent Nova2OE epistasis experiment. B.W., L.A.I, N.A.G and N.Y. analyzed and interpreted the results. R.D and Y.Y. provided the Nova antibody and conditional mouse lines to remove *Nova1* and *Nova2*. J.D. assisted with designing plasmids and viruses. Q. X and L.G. produced viruses. B.W., L.A.I, and G.F. wrote the paper with review and editing provided by all authors.

---

**Figure S1:**
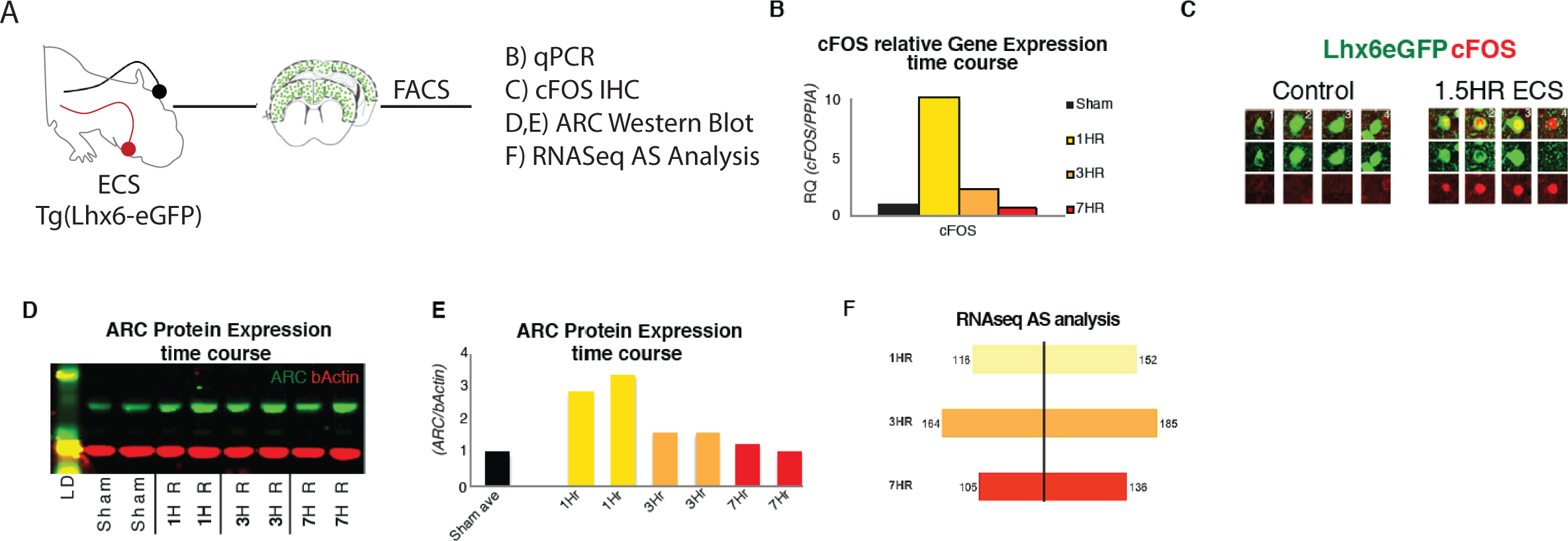
Acute increases in neuronal activity induces immediate early gene expression and differential splicing within SST+ cINs in vivo (Related to Figure 2)|. A) Schematic of experimental approach: P8 tgLhx6eGFP animals were subjected to ECS (left)’ then following a time course of 1 Hr, 3 hr or 7 hrs the S1 cortex was dissected (middle) and GFP+cINs were isolated by FACS for qPCR, Western blot, and RNAseq analysis and IHC (right). B) Quantification of relative mRNA expression (RQ) of cFOS (normalized to housekeeping gene PPIA) in ctl/sham animals (black), 1hr (yellow), 3hrs (orange), and 7hrs (red) following ECS within cINs. C) Immunostaining of cFOS in sham (red) and eGFP vs ECS treated animals, showing the expression of cFOS 1.5 hours post ECS activity induction. D) Representative western blot of ARC protein expression within sham treated (two replicates), 1hr (two replicates), 3hrs (two replicates), and 7hrs (two replicates) following ECS within cINs. E) Fold of ARC protein expression (normalized to b-actin) in ctl/sham treated (black), 1hr (yellow), 3hrs (orange), and 7hrs (red) following ECS within cINs. F) Magnitude of differential alternative splicing events from the comparison of sham cINs to cINs 1hr (yellow, 268 events), 3hrs (orange, 349 events), and 7 hrs (red, 241 events) following ECS.

**Figure S2:**
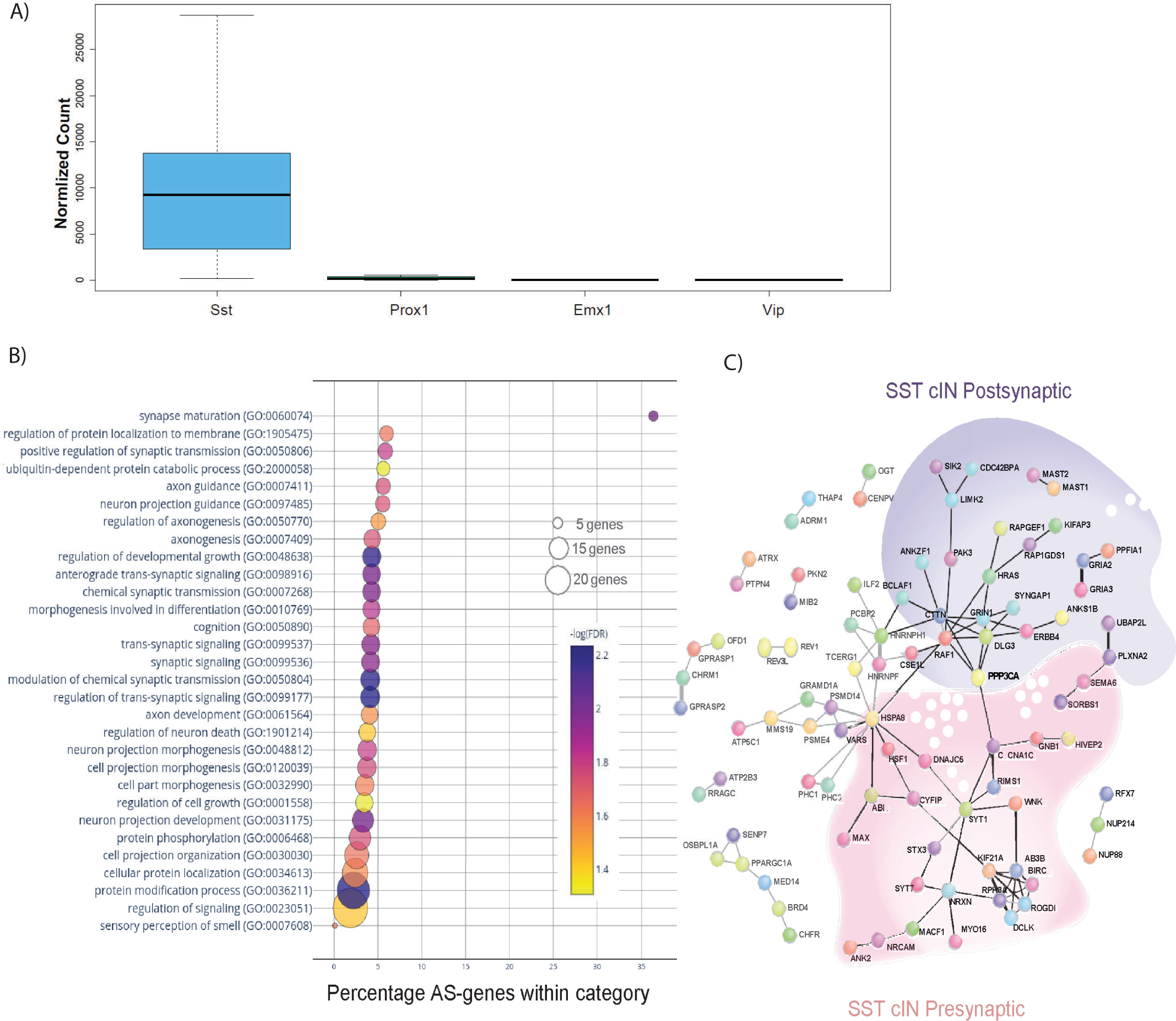
Neuronal activity influences alternative splicing and Nova expression within SST cINs (Related to Figure2) A) Normalized counts of SST, Prox1, Emx1 and Vip gene expression from the RNA sequencing analysis from control tissue indicating the purity of our sample preparation to be specific for SST+ cINs. B) Bubble dot plot of gene ontology (GO) most significant terms for the genes subjected to activity-dependent alternative splicing within SST+ cINs (false discovery rate (FDR)<0.05), x-axis is the enrichment of the activity-dependent AS genes in the GO catagory (# of genes in GO category from SST transcriptome/ # of genes activity-dependent AS in category). Color of dot indicates magnitude of significance (-log10 transform FDR, none shown above FDR <0.05) and size corresponds to number of genes in catagory. C) Protein-protein interaction (PPI) network formed from 312 activity-depdendent spliced genes in SST cINs with Disease Association Protein-Protein Link Evaluator (DAPPLE) (Rossin et al., 2011) and performed over 10,000 permutations (pVal<0.00009). Green shading-post-synaptic gene network, pink shading-pre-synaptic gene network.

**Figure S3:**
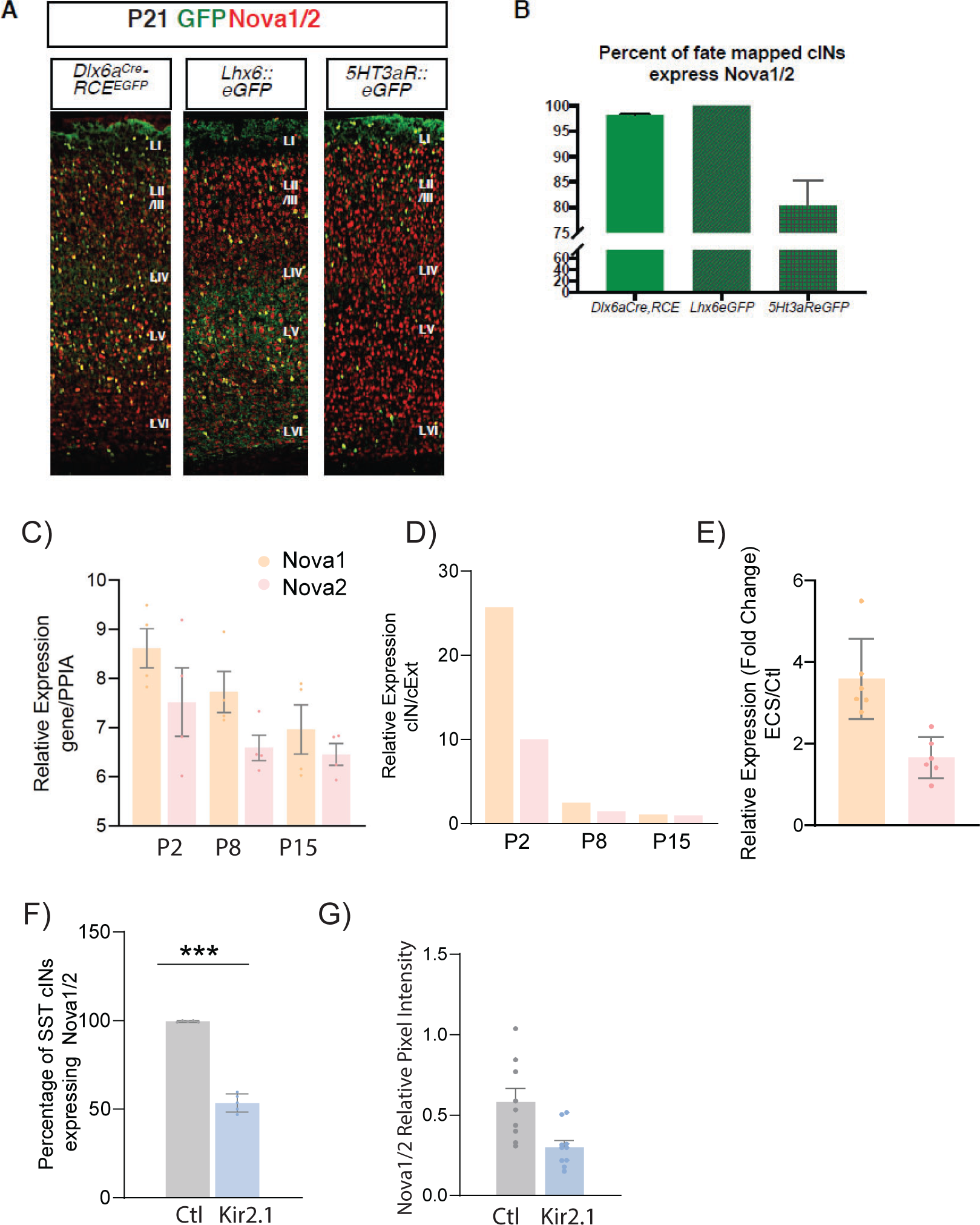
Nova1/2 AS factors expression within the MGE and cINs (Related to Figure 3) A) Representative IHC image of brain section at P21 from Dlx6aCre;RCEeGFP, Lhx6::eGFP and 5HT3aR::eGFP. Anti-Nova1/2 (red); GFP (green). B) Quantification of eGFP cells expressing Nova1/2. 100% of Lhx6::eGFP cells express Nova1/2. C) Relative gene expression of Nova1 (orange) and Nova2 (pink), normalized to house-keeping gene Peptidyl prolyl isomerase A (PPIA) using qPCR from Lhx6-eGFP sorted cINs at Postnatal age (P) P2, P8 and P15 (n=4 mice each, S1 cortex only). D) Fold change of the relative expression of Nova1 and Nova2 between cINs and excitatory neurons (cExt) showing an enrichment of Nova expression in cINs at early developmental ages (n=4 mice each, S1 cortex only). E) Relative expression of Nova1 and Nova2 genes (using qPCR) of ECS induced SST cINs relative to controls (n=4-6 mice, S1 cortex only; **pVal=0.002, Nova1; **pVal=0.005, Nova2). F) Quantification of the number of Nova1/2-expressing SST+ cINs of control AAV2/1-Flex-mCherry (grey) and AAV2/1-Flex-Kir2.1-P2A-mCherry (blue) injected animals. (n=7, S1 cortex, ∼27 cells each; ***pVal= 0.0001). G) Representative images of Nova1/2 expression, Left: control SST cIN (injected with mCherry), Right: KIR2.1+ SSt cIN at P21. Right, Quantification of Nova1/2 protein pixel intensity (normalized to area) from ctl SST cINs (grey) and KIR2.1+ SST cINs (blue) (n=10; **pVal=0.006).

**Figure S4:**
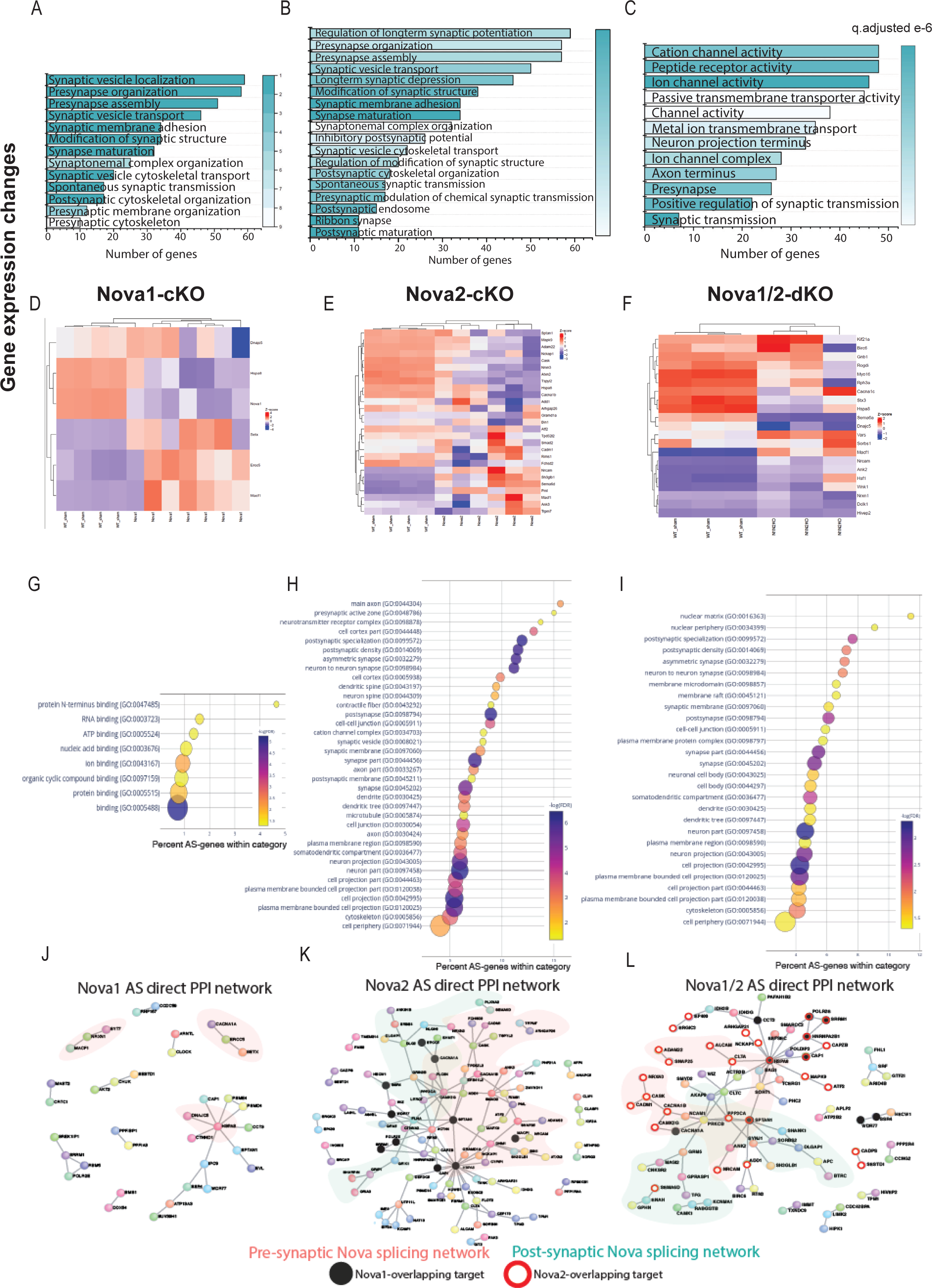
Nova2 controls the majority of gene expression and splicing events of the Nova1/2 family within SST+ cINs and these events coalese into GO categories and PPI networks related to pre and post-synaptic development of SST cINs. A-C) Gene ontology analysis of differentially expressed synaptic genes in Nova1-cKO (A), Nova2-cKO (B), and Nova1/2-dKO (C). Color bar indicates adjusted qvalue. D-F) Examples of some upregulated and downregulated genes in Nova1-cKO (D), Nova2-cKO (E) and Nova1/2-dKO (F). G) Bubble dot plot of most significant GO terms for the genes subjected to AS within SST-Nova1-cKO, x-axis is the percent enrichment of the AS genes in the GO category (#genes SST-Nova1 AS in category divided by #genes in GO category from SST transcriptome). Color of dot indicates magnitude of significance (-log10 FDR, none shown above FDR<0.05) and size corresponds to number of genes in category. H) Bubble plot of most significant GO terms of SST_Nova2-cKO genes subjected to AS illustrating the substantial enrichment of Nova2 dependent events to synaptic development. I) Same as G-H but for Nova1/2-dKO J) Protein-protein interaction (PPI) network formed from 124 Nova1-cKO spliced genes in SST cINs with DAPPLE (10,000 permutations, pVal<0.09), pink shading labels genes that belong in synapse related GO categories. K) PPI network formed from 339 Nova2-cKO spliced genes in SST-cINs with DAPPLE (10,000 permutations, pVal<0.00009). Black dots indicate shared genes with Nova1-cKO, green shading labels postsynaptic genes that belong in synapse related GO categories, pink shading labels pre-synaptic genes in synapse GO categories. L) PPI network in Nova1/2-dKO. 270 Nova1/2-dKO splice genes in SST cINs with DAPPLE (pVal<0.00009) Red dots indicate overlap with Nova2, whereas black dots indicate overlap with Nova1-cKO.

**Figure S5:**
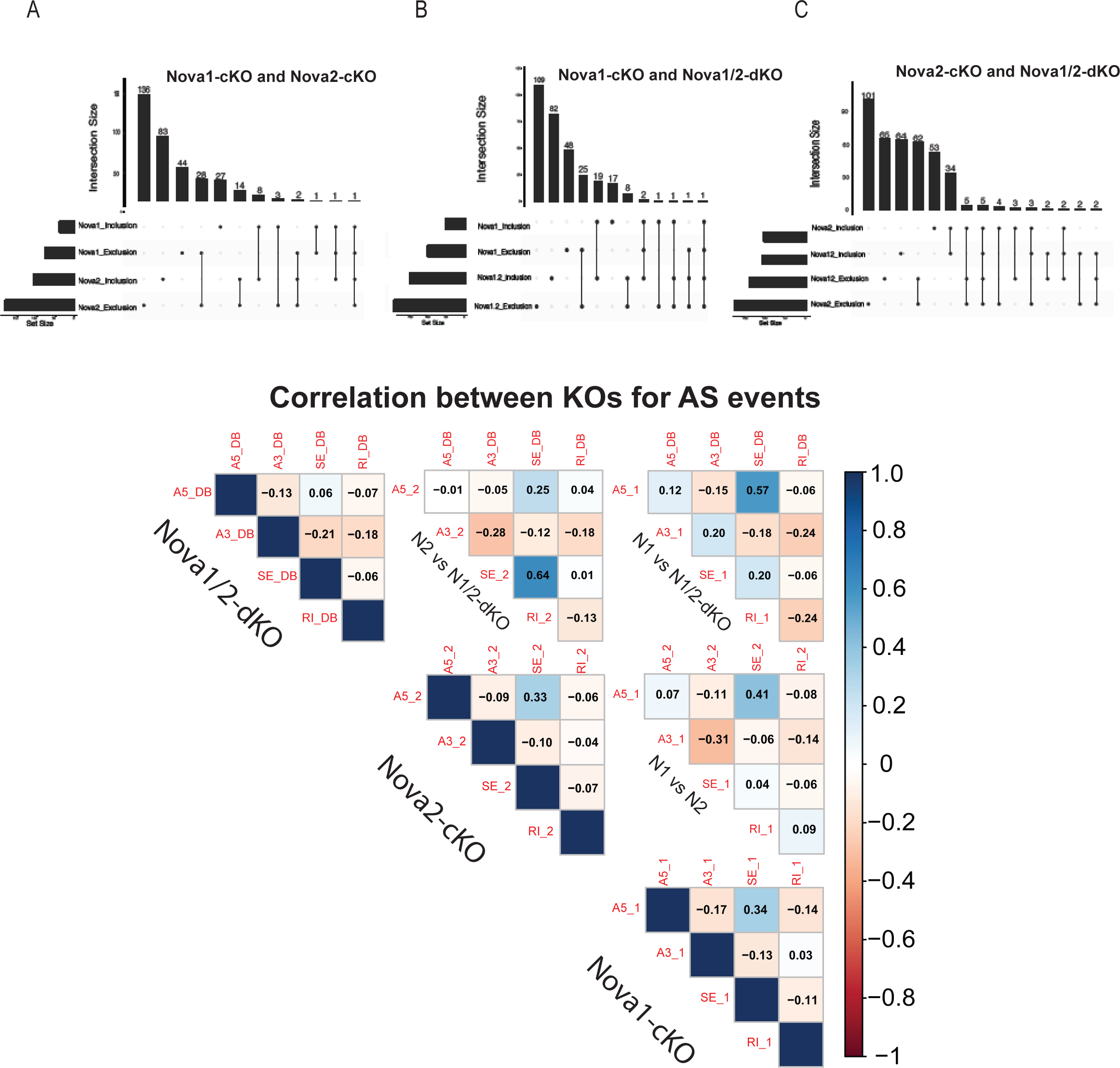
Overlap in Alternative Splice events (Related to Figure 4) A) Quantification of the overlap of SST-Nova1-cKO and Nova2-cKO splicing events. Bottom left horizontal bars indicate the number of genes subjected to AS events for each data set: Nova1_inclusion splice events (43), Nova1_exclusion splicing events (81), Nova2_inclusion splicing events (122), Nova2_exclusion splicing events (217). Top histogram bars indicate the magnitude of overlap between the data sets indicated by a filled black circle below (i.e. 136 of Nova2_exclusion events overlap with the other sets, 28 of Nova1_exclusion events overlap with Nova2_exclusion etc.) B) Similar to A but Nova1-cKO and Nova1/2-dKO overlap. Bottom left horizontal bars indicate number of genes subjected to AS events for each dataset. Nova1_inclsion splicing events (43), Nova1_exclusion (81). Nova1/2_inclusion (108), Nova1/2_exclusion (162). Top histogram bars indicate the magnitude of overlap between the two datasets. 25 of Nova1_exclusion events overlap with Nova1/2_exclusion; 19 of Nova1_inclusion overlaps with Nova1/2_inclusion etc. C) Similar to B, but Nova2-cKO and Nova1/2-dKO overlap. 62 of Nova2_exclusion events overlap with Nov1/2; 34 of Nova2_inclusion events overlap with Nova1/2. D) Overall correlation between Nova1, Nova2 and Nova1/2-dKO for each type of splice event.

**Figure S6:**
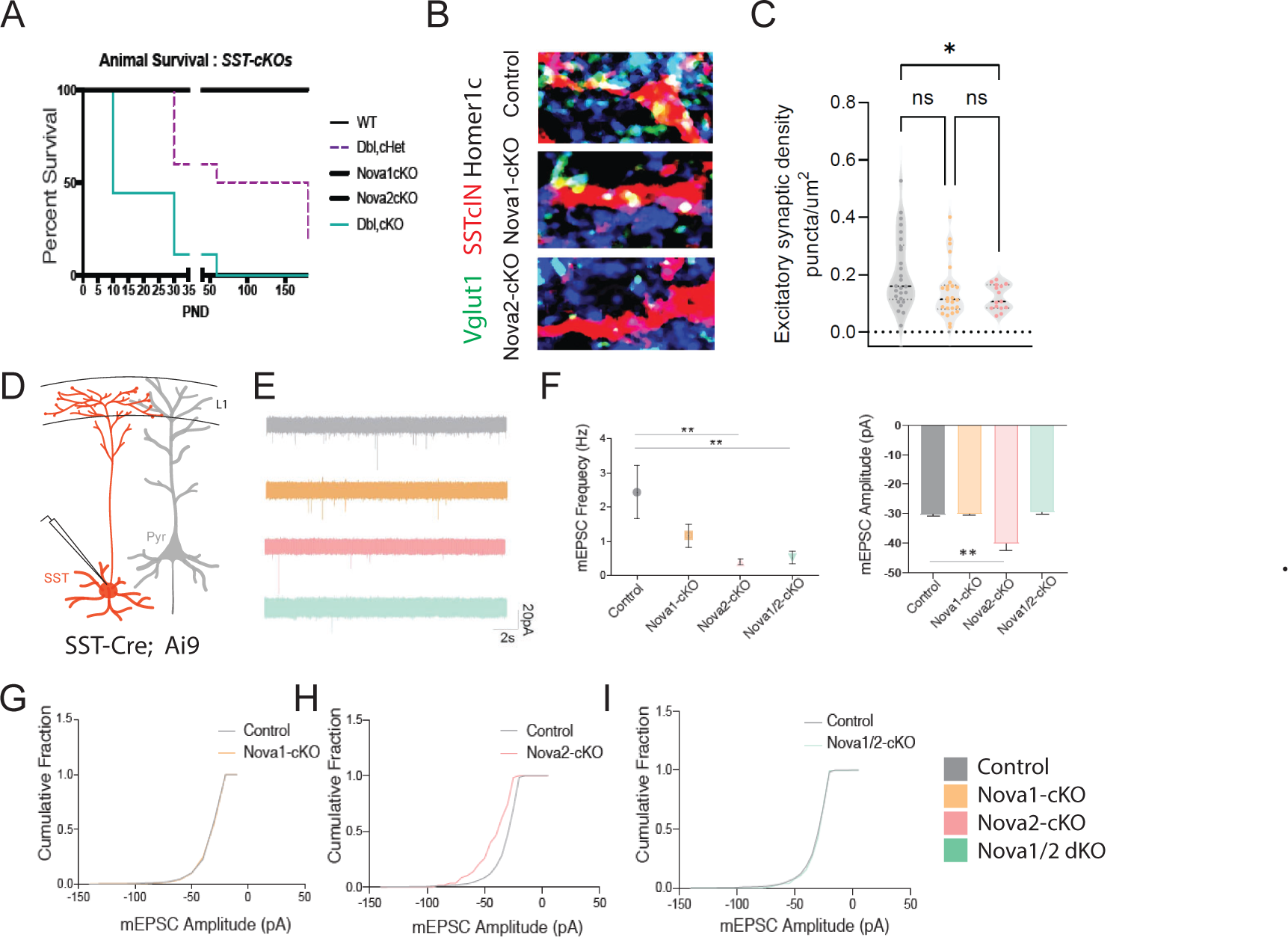
Conditional loss of Nova1/2 within SST+ cINs impacts animal survival and disrupts their afferent synaptic connectivity (Related to Figure 5) A) Survival plot of conditional knockouts within SSTCre;Ai9 animals (WT SSTCre; Ai9 animals black line, double heterozygous (het) SSTCre; Ai9, Nova1f/+, Nova2f/+ slashed purple line (Dcl,cHet), SSTCre; Ai9, Nova1f/f black line, SSTCre; Ai9, Nova2f/f black line, double SSTCre;t Ai9, Nova1f/f Nova2f/f turquoise line) at P8 50% of double conditional SSTCre;t Ai9, Nova1f/f Nova2f/f (Dbl,cKO) animals have deceased by P50 90% of double conditional animals have deceased. Whereas, at P30 40% of Dbl, cHet animals and by P60 50% have deceased. WT and singular conditional mutants do not exhibit decreased survival. B) SST+ afferents: IHC of representative SST+ cIN dendrite of anti-RFP (red), anti-VGLUT1 (green), and anti-Homer1c (blue) to label excitatory synaptic puncta overlapping with SST+ cINs dendrites (RFP+/VGLUT1+/Homer1c+ puncta, white) in S1 cortex of SST-ctl, SST-Nova1 and SST-Nova2 mutant animals. C) Quantification of the density of excitatory afferent synapses onto SST+ cINs within L2/3 and L5/6 of S1 cortex of SST-ctl, SST-Nova1 and SST-Nova2 mutant animals. (n=23, 3 mice each; *pVal=0.028, SST-Nova1; *pVal=0.012, SST-Nova2). D) Schematic of experimental approach recording mini-excitatory postsynaptic potentials (mEPSCs) in SST+ cINs in the S1 cortex (red). H) Quantification of mEPSCs frequencies from SST+ cIN Ctl,SST-Nova1 and SST-Nova2 mutant animals (n=15 cells from 3 mice of each genotype) E) Representative mEPSCs recordings from SST cINs from: Top to bottom: wt animals, grey traces (SSTCre; Ai9 or SSTCre; Ai9, Nova1f/+ or SSTCre; Ai9, Nova2f/+), Nova1-cKO animals, orange traces (SSTCre;t Ai9, Nova1f/f), Nova2-cKO animals, pink traces (SSTCre; Ai9 Nova2f/f), and double Nova1/2-cKO, turquoise traces (SSTCre; Ai9, Nova1f/f Nova2f/f). Scale bar:20pA and 2 seconds. F) Quantification of mEPSC frequencies and amplitude recorded from SST cINs in control animals, grey dot (SSTCre; Ai9 or SSTCre; Ai9, Nova1f/+ or SSTCre; Ai9, Nova2f/+), Nova1-cKO animals, orange square (SSTCre; Ai9, Nova1f/f), Nova2-cKO animals, pink triange (SSTCre; Ai9, Nova2f/f), and double Nova1/2-cKO, turquoise upside-down triangle. **pVal= <0.005 for wt vs. Nova2-cKO and wt vs. Nova1/2-cKO, G) Cumulative probablility distributions of mEPSC amplitudes from recordings of wt SST cINs, grey line, and Nova1-cKO SST cINs, orange line, exhibiting no difference. H) Cumulative probablility distributions of mEPSC amplitudes from recordings of wt SST cINs, grey line, and Nova2-cKO SST cINs, pink line, exhibiting a significant increase in the amplitude of mEPSCs in Nova2-cKO SST cINs. I) Cumulative probablility distributions of mEPSC amplitudes from recordings of wt SST cINs, grey line, and Nova1/2-cKO SST cINs, turquoise line, exhibiting no difference.

**Figure S7:**
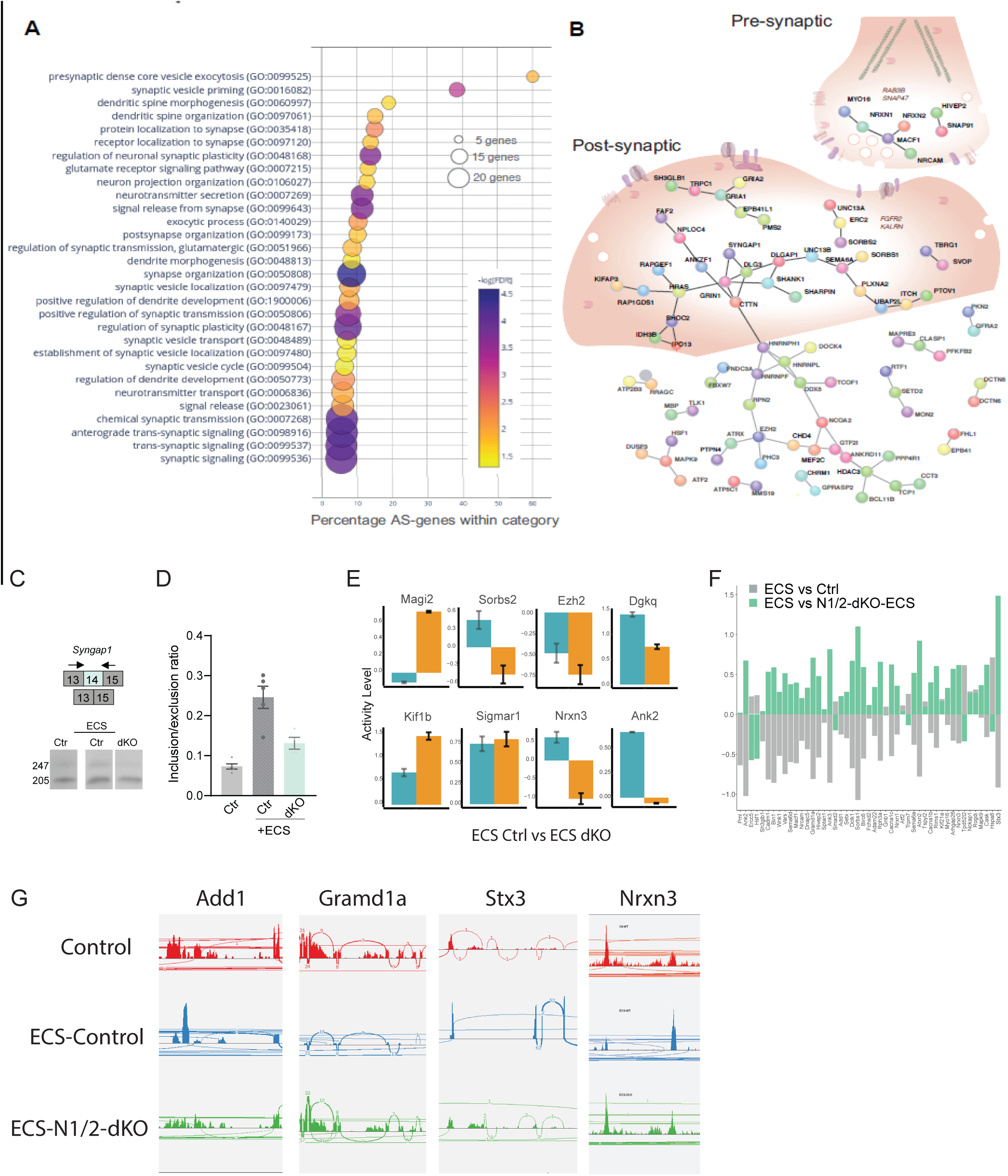
Nova RNA binding proteins control activity dependent AS in SST cINs during development (Related to Figure 6) A) Bubble dot plot of the most significant GO terms for the genes undergoing Nova1/2 activity dependent AS splicing within SST+ cINs (all GO terms shown FDR<0.05). B) Schematic of a SST+cIN presynaptic inhibitory axonal puncta (top right) and a SST+ cIN excitatory post-synaptic density (middle) overlaid on top of the significant DAPPLE generated PPI direct network from the 356 genes undergoing Nova1/2 dependent activity induced AS (***pVal=0.00009, 10,1000 permutations). C) Example RT-PCR validation of alternative splicing (AS) events of activity- and Nova1/2-dependent alternative exon usage within the gene Syngap1 (top), bottom, Gel image of RT-PCR product from the amplification of exon13 to exon 15 within SST-ctl cINs (Ctl) (left), ECS-treated Ctl (middle), and ECS-treated SST-dKO (right). D) Quantification of RT-PCR AS events of Syngap1. **pVal=0.0001 Ctl vs Ctl+ECS; **pVal=0.004 Ctl+ECS vs SST-dKO+ECS. E) Examples of genes with both GE and AS changes. y-axis represents activity level (FC and pValue) of either GE (teal) or AS (orange). F) Gene expression changes observed in ECS vs control are partially abolished by Nova1/2-dKO G) Alternatice splicing changes due to activity induction (ECS) are significantly abolished in the N1/2-dKO (with ECS). A few example genes are presented showing the activity dependent exclusion/inclusion of certain exons are no longer present in the dKO Red: Control; Blue: ECS; Green: N1/2-dKO+ECS

**Figure S8:**
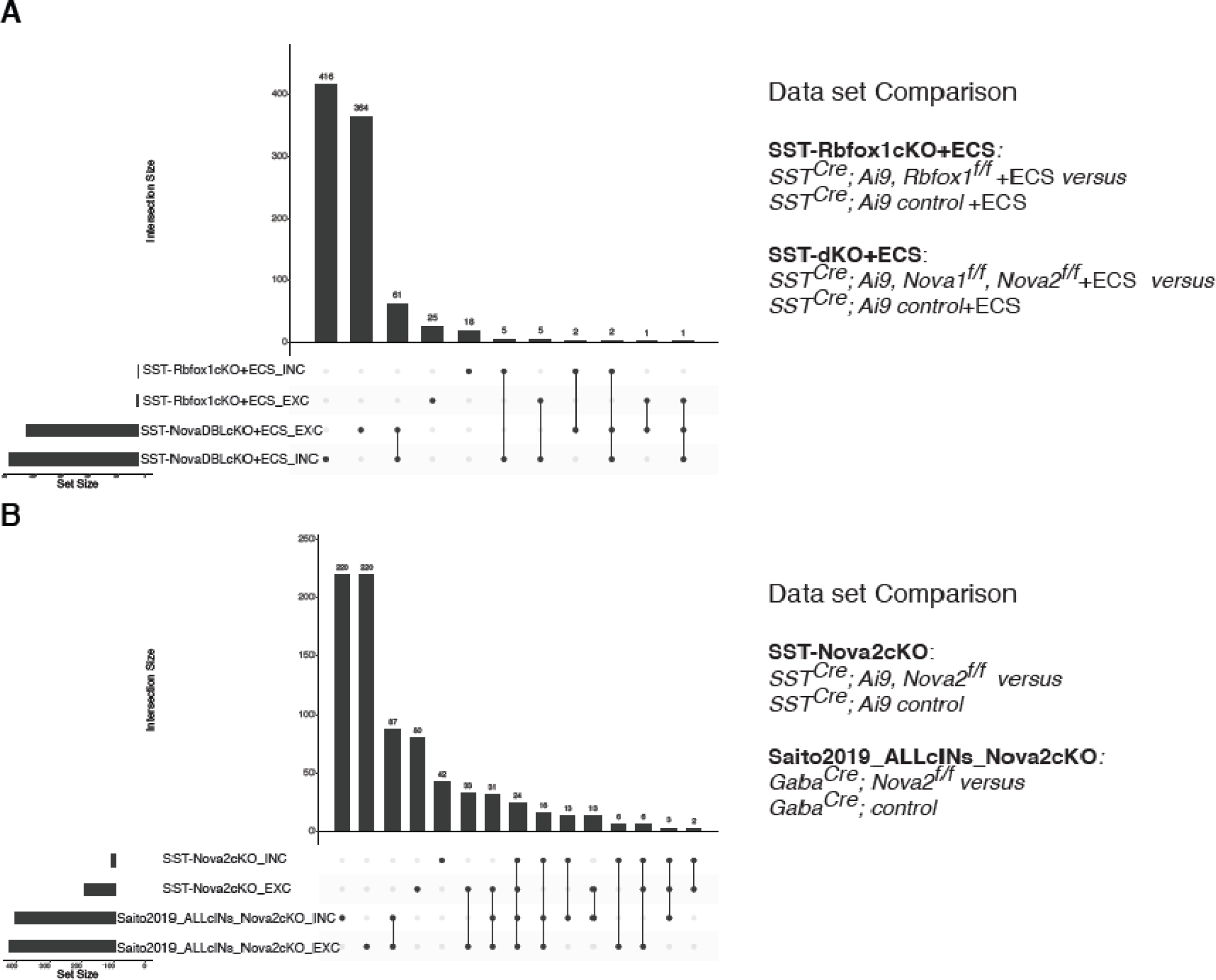
Nova1/2 controls the activity dependent splicing of large and unique pool of mRNAs compared to Rbfox1 within SST cINs and SST specific Nova2 AS genes overlap well with pan-cIN Nova2 AS genes. A) Quantification of the overlap of SST-Rbfox-cKO + ECS (differential splicing events from the comparison of SST-cIN+ECS to SST-RbfoxcKO+ECS) and SST-dKO (differential splicing events from the comparison of SST cINS +ECS to SST-Nova1/2-dKO+ECS). Bottom left horizontal bars indicate the number of genes subjected to AS events for each data set. Top vertical bars indicate the number of overlapping genes corresponding to the black dot below indicating the data set identity. B) Qyantification of the overlap of SST-Nova2 cIN splicing events (differential splicing events from the comparison of Ctrl SST cINs to SST-Nova2-cKO) with the dataset generated by Saito et al 2019 utilizing a mouse cross GadCre and Nova2f/f (differential splicing events from Ctrl cINs vs cINs-Nova2-cKO).

## Notes

https://github.com/IbrahimLab-23/Nova-proteins-and-synaptic-integration-of-Sst-interneurons

### Competing Interest Statement

Gord Fishell and Jordane Dimidschtein are founders of Regel therapeutics. All other authors declare no competing interest.

### Summary of Updates

New data analyzed and added. Figure 7 is revised. author affiliation updated.

