## Supplementary Tables for "Nova proteins direct synaptic integration of somatostatin interneurons through activity-dependent alternative splicing"

| | *Control* | *Nova1-cKO* | *Nova2-cKO* | *N1/N2 dKO* | *One-Way ANOVA p-value* |
|---|---|---|---|---|---|
| Input resistance [MΩ] | 213±10 | 192 ± 15 | 250±18 | 250 ± 20 | 0.03 |
| Sag ratio | 1.05 ± 0.033 | 1.087 ± 0.037 | 1.077± 0.037 | 1.1±0.05 | 0.07 |
| Rheobase [pA] | 120± 25 | 83 ± 15 | 25±3 | 128±30 | 0.01 |
| Peak amplitude (First Spike) [mV] | 86±7 | 81±3 | 87±4 | 60±9 | 0.007 |
| AHP (First Spike) [mV] | -9.6 ± 4 | -9.7 ± 0.85 | -6±1 | -8.6 ± 1.1 | 0.41 |
| Halfwidth (First Spike) [ms] | 1.3 ± 0.3 | 1.6 ± 0.4 | 1.9±0.3 | 2.5 ± 0.24 | 0.002 |
| Threshold (First Spike) [mV] | -34.29 ± 3.08 | -42.24 ± 5.93 | -36.6±3.9 | -24.1 ± 4.5 | 0.0013 |

Table S1

| | Gene Symbol | Ctl_WT_Exon Coverage | ECSWT_Exon Coverage | ECS2KO_Exon Coverage | Total # Exon | WT_IncLevel | ECS_IncLevel | IncLevelDifference | NDBKO_IncLevel | ECS_IncLevel | IncLevelDifference |
|---|---|---|---|---|---|---|---|---|---|---|---|
| 1 | Macf1 | 99 | 305 | 97 | 102 | "0.809,0.84" | "0.977,1.0" | -0.122 | "0.337,0.179" | "0.232,0.002" | 0.113 |
| 2 | Nrxn1 | 44 | 90 | 46 | 24 | "0.351,0.758" | "0.061,0.068" | 0.469 | "0.916,0.359" | "0.061,0.068" | 0.588 |
| 3 | Nrcam | 73 | 306 | 62 | 35 | "0.571,0.257" | "0.2,0.016" | 0.29 | "0.583,0.477" | "1.0,0.958" | -0.327 |
| 4 | Sema6a | 126 | 184 | 95 | 20 | "0.622,1.0" | "0.215,0.014" | 0.468 | "0.0,0.39" | "0.215,0.014" | 0.109 |
| 5 | Sorbs1 | 66 | 202 | 66 | 33 | "0.816,0.876" | "0.428,0.009" | 0.508 | "0.913,0.629" | "0.159,0.329" | 0.418 |
| 6 | Hivep2 | 165 | 362 | 137 | 13 | "0.432,1.0" | "0.12,0.0" | 0.507 | "0.008,0.165" | "0.241,0.328" | -0.158 |
| 7 | Wnk1 | 57 | 197 | 39 | 34 | "0.673,0.567" | "0.402,0.0" | 0.433 | "0.71,0.494" | "0.402,0.0" | 0.25 |
| 8 | Kif21a | 70 | 193 | 80 | 38 | "0.893,0.853" | "0.581,0.239" | 0.457 | "0.264,0.577" | "0.511,0.0" | 0.243 |
| 9 | Birc6 | 107 | 138 | 83 | 73 | "0.182,0.078" | "0.115,0.003" | 0.21 | "0.461,0.131" | "0.158,0.879" | -0.34 |
| 10 | Rogdi | 88 | 118 | 53 | 16 | "0.028,0.0" | "0.025,0.298" | -0.148 | "0.047,0.032" | "0.025,0.298" | -0.132 |
| 11 | Myo16 | 80 | 200 | 72 | 36 | "1.0,1.0" | "1.0,0.435" | 0.282 | "1.0,0.987" | "1.0,0.435" | 0.279 |
| 12 | Ank2 | 206 | 784 | 246 | 52 | "0.278,0.334" | "0.491,0.532" | -0.185 | "0.705,0.416" | "0.491,0.532" | 0.18 |

**Table S2.1**

| | Gene Symbol | Ctl_WT_Exon Coverage | ECSWT_Exon Coverage | ECS2KO_Exon Coverage | Total # Exon |
|---|---|---|---|---|---|
| 1 | Macf1 | 99 | 305 | 97 | 102 |
| 2 | Ercc5 | 10 | 19 | 12 | 15 |
| 3 | Setx | 13 | 30 | 10 | 28 |
| 4 | Dnajc5 | 67 | 186 | 51 | 7 |
| 5 | Hspa8 | 1885 | 1274 | 1488 | 9 |
| 6 | Sema6d | 41 | 101 | 53 | 23 |
| 7 | Trpm7 | 12 | 20 | 10 | 40 |
| 8 | Arhgap26 | 16 | 61 | 14 | 27 |
| 9 | Fchsd2 | 33 | 78 | 39 | 21 |
| 10 | Cadm1 | 93 | 182 | 145 | 14 |
| 11 | Cask | 121 | 334 | 75 | 28 |
| 12 | Tspyl2 | 54 | 95 | 36 | 7 |
| 13 | Nrxn1 | 44 | 90 | 46 | 24 |
| 14 | Tpd52l2 | 78 | 65 | 57 | 9 |
| 15 | Ank3 | 170 | 246 | 129 | 53 |
| 16 | Cacna1b | 89 | 162 | 36 | 49 |
| 17 | Add1 | 19 | 178 | 23 | 18 |
| 18 | Rims1 | 70 | 87 | 45 | 34 |
| 19 | Smad2 | 27 | 50 | 50 | 11 |
| 20 | Pml | 10 | 29 | 19 | 11 |
| 21 | Atf2 | 60 | 127 | 60 | 16 |
| 22 | Sptan1 | 239 | 364 | 215 | 59 |
| 23 | Mapk9 | 137 | 118 | 72 | 13 |
| 24 | Adam22 | 178 | 345 | 178 | 34 |
| 25 | Nrcam | 73 | 306 | 62 | 35 |
| 26 | Gramd1a | 51 | 327 | 57 | 21 |
| 27 | Nckap1 | 161 | 209 | 102 | 32 |
| 28 | Sh3glb1 | 113 | 192 | 149 | 12 |
| 29 | Atxn2 | 91 | 300 | 71 | 25 |
| 30 | Bin1 | 104 | 447 | 95 | 20 |
| 31 | Sema6a | 126 | 184 | 95 | 20 |
| 32 | Sorbs1 | 66 | 202 | 66 | 33 |
| 33 | Hivep2 | 165 | 362 | 137 | 13 |
| 34 | Gnb1 | 1376 | 945 | 1254 | 12 |
| 35 | Cacna1c | 46 | 158 | 34 | 50 |
| 36 | Wnk1 | 57 | 197 | 39 | 34 |
| 37 | Kif21a | 70 | 193 | 80 | 38 |
| 38 | Birc6 | 107 | 138 | 83 | 73 |
| 39 | Rph3a | 401 | 774 | 370 | 23 |
| 40 | Rogdi | 88 | 118 | 53 | 16 |
| 41 | Dclk1 | 319 | 1228 | 252 | 18 |
| 42 | Myo16 | 80 | 200 | 72 | 36 |

| | | | | | |
|---|---|---|---|---|---|
| 43 | Nrxn3 | 53 | 83 | 37 | 24 |
| 44 | Ank2 | 206 | 784 | 246 | 52 |
| 45 | Stx3 | 64 | 102 | 50 | 12 |
| 46 | Hsf1 | 27 | 193 | 25 | 17 |
| 47 | Vars | 16 | 52 | 19 | 30 |

**Table S2.2**
